# Low acetylcholine during early sleep is important for motor memory consolidation

**DOI:** 10.1101/494351

**Authors:** Samsoon Inayat, Qandeel, Mojtaba Nazariahangarkolaee, Surjeet Singh, Bruce L. McNaughton, Ian Q. Whishaw, Majid H. Mohajerani

**Affiliations:** 1Canadian Centre for Behavioural Neuroscience, University of Lethbridge, Lethbridge, Alberta, Canada; Center for the Neurobiology of Learning and Memory, University of California, Irvine, California, USA

**Keywords:** Rotarod, skilled reach task, physostigmine, REM/NREM sleep, DREADD

## Abstract

The synaptic homeostasis theory of sleep proposes that low neurotransmitter activity in sleep is optimal for memory consolidation. We tested this theory by asking whether increasing acetylcholine levels during early sleep would disrupt motor memory consolidation. We trained separate groups of adult mice on the rotarod walking and skilled reaching for food tasks, and after training, administered physostigmine, an acetylcholinesterase inhibitor, to increase cholinergic tone in subsequent sleep. Post-sleep testing suggested that physostigmine impaired motor skill acquisition. Home-cage video monitoring and electrophysiology revealed that physostigmine disrupted sleep structure, delayed non-rapid-eye-movement sleep onset, and reduced slow-wave power in the hippocampus and cortex. The impaired motor performance with physostigmine, however, was not solely due to its effects on sleep structure, as one hour of sleep deprivation after training did not impair rotarod performance. A reduction in cholinergic tone by inactivation of cholinergic neurons during early sleep also affected rotarod performance. Administration of agonists and antagonists of muscarinic and nicotinic acetylcholine receptors revealed that activation of muscarinic receptors during early sleep impaired rotarod performance. The experiments suggest that the increased slow wave activity and inactivation of muscarinic receptors during early sleep due to reduced acetylcholine contribute to motor memory consolidation.

---

Memory formation consists of a number of processes; encoding which involves stimulus-induced gene-expression within cells and consolidation which happens through formation and strengthening of synaptic connections (Ebbinghaus H 1885-Republished Translation 1964; Jenkins JG and KM Dallenbach 1924; McGaugh JL 2000; Dudai Y 2004; Frankland PW and B Bontempi 2005; Diekelmann S and J Born 2010; Clopath C 2012; Rasch and J Born 2013; Yang G et al. 2014; Dudai Y et al. 2015; Chambers AM 2017). Memory consolidation can happen in both awake and sleep states, can last from seconds to years, and might involve different processes for different kinds of memory (Clopath C 2012; Rasch and J Born 2013; Chambers AM 2017), with procedural memory and episodic memories consolidated during rapid eye movement (REM) or non-REM (NREM) phases/stages of sleep respectively (Yaroush R et al. 1971; Plihal W and J Born 1997), or through cyclic succession of NREM and REM (Giuditta A et al. 1995; Ambrosini MV and A Giuditta 2001; Genzel L et al. 2014; Boyce R et al. 2016). Reactivation of memories during sleep (Wilson MA and BL McNaughton 1994; Kudrimoti HS et al. 1999; Sutherland GR and B McNaughton 2000; Kali S and P Dayan 2004; Euston DR et al. 2007; Buhry L et al. 2011; Tononi G and C Cirelli 2014; Yang G *et al*. 2014; Klinzing JG et al. 2018; Olcese U et al. 2018), formation and strengthening of spines during NREM and REM phases of sleep, and synapse pruning during REM sleep might also play a role (Yang G *et al*. 2014; Li W et al. 2017). The synaptic homeostasis theory of memory consolidation suggests a mechanism through which synaptic strengthening could occur. It proposes that low levels of neuromodulators, such as acetylcholine, dopamine, serotonin, during NREM sleep permit overall synaptic downscaling while allowing recently activated synapses to be relatively more active. Thus, improved signal-to-noise provides a mechanism for synaptic stabilization, memory replay, and memory consolidation (Tononi G and C Cirelli 2003; Vyazovskiy VV et al. 2009; Tononi G and C Cirelli 2014).

Consistent with the synaptic homeostasis theory, low levels of acetylcholine (ACh) allow synaptic pruning/formation and episodic memory consolidation during NREM sleep (Hasselmo ME 1999; Walker MP 2009; Alger SE et al. 2015; Miyamoto D et al. 2017). Whether motor learning is similarly enhanced by low ACh levels is uncertain. Learning of motor tasks is considered a form of procedural learning that involves the acquisition of skills and sequences of movements such as those involved in rotarod walking and skilled reaching for food. Studies on the effects of lesions that lower cortical ACh on motor learning have produced mixed results (Conner, Culberson et al. 2003; Gharbawie and Whishaw 2003). Other evidence suggests that increasing cholinergic tone in humans during post-learning NREM does not affect consolidation of procedural memories (Gais S and J Born 2004). Nevertheless, disrupting cholinergic levels through sleep deprivation also affects motor memory. Specifically, reactivation of motor task-specific neurons during NREM sleep is involved in forming new synapses after motor learning in mice but consolidation is impaired in sleep deprived animals (Yang G *et al*. 2014). Furthermore, there is a proposed relationship between the amount of slow-wave activity (SWA) during NREM sleep and motor learning. For example, humans learning a reaching task display local increases in SWA in the parietal cortex (Landsness EC et al. 2011) whereas arm immobilization induces a local decrease in SWA (Huber R et al. 2006). SWA also significantly improves visual texture discrimination skills (Gais S et al. 2000). In mice, performance of a skilled forelimb reaching for food task increases local SWA in motor cortex (Hanlon EC et al. 2009). In sum, this evidence points to a role for decreased cholinergic tone during NREM sleep in motor memory consolidation, but a more definitive conclusion requires a direct test of the idea.

Here, we performed a direct test of the role of cholinergic activity during early NREM sleep in the consolidation of motor memory with mice performing a rotarod task and a skilled reaching task. The effects of acetylcholine levels on sleep structure and phases were then manipulated. We also related acetylcholine levels in post-training sleep to subsequent motor performance and we investigated the role of muscarinic and nicotinic acetylcholine receptors in motor memory consolidation. In sum, we propose that low acetylcholine level during NREM sleep contributes to the consolidation of motor memory.

## MATERIALS AND METHODS

All experiments were performed in accordance with Canadian Council of Animal Care and were approved by the University of Lethbridge Animal Welfare Committee.

### Experimental Animals

Adult wild-type (WT) mice (C57 Bl/6J, Jackson laboratories), N=109, and transgenic mice (Chat-Cre::CAG-hM4Di), N=18, 3-6 months old, both male and female, and weighing 20-30g were used in this study. Transgenic mice had inhibitory (hM4Di) DREADDs in cholinergic neurons, which were activated by the inert molecule clozapine-*N*-oxide (CNO) to reduce release of acetylcholine. For generating these mice, CAG-hM4Di mice (B6N.129-Gt(ROSA)26Sortm1(CAG-CHRM4*,-mCitrine) Ute/J) were crossed with Chat-Cre mice (B6;129S6-Chattm2(cre)Lowl/J) to enable expression of CAG hM4Di in cholinergic neurons of the brain (Stachniak TJ et al. 2014; Vardy E et al. 2015). All mice were housed up to four per cage and were provided food and water ad libitum. They were kept in a controlled temperature (22°C), humidity, and light with a 12:12 light/dark cycle (lights on at 7:30am). All testing and training was performed during the light phase of the cycle at the same time each day.

### Motor Learning Tasks

#### Rotarod Task

In the rotarod task, mice learn to balance while walking on an elevated rotating drum whose speed is gradually increasing (Fig. 1A). As they learn to keep their balance, they can stay on the rotating drum for longer times at higher drum rotational speeds. Eighty four C57/BL6 and 17 Chat-Cre::CAG-hM4Dimice were used and were randomly divided into control and experimental groups. The number of males and females in individual groups are mentioned below. However, because of the constraints on the availability of mice, the number of males and females are not always equal in individual groups.

**Figure 1.**
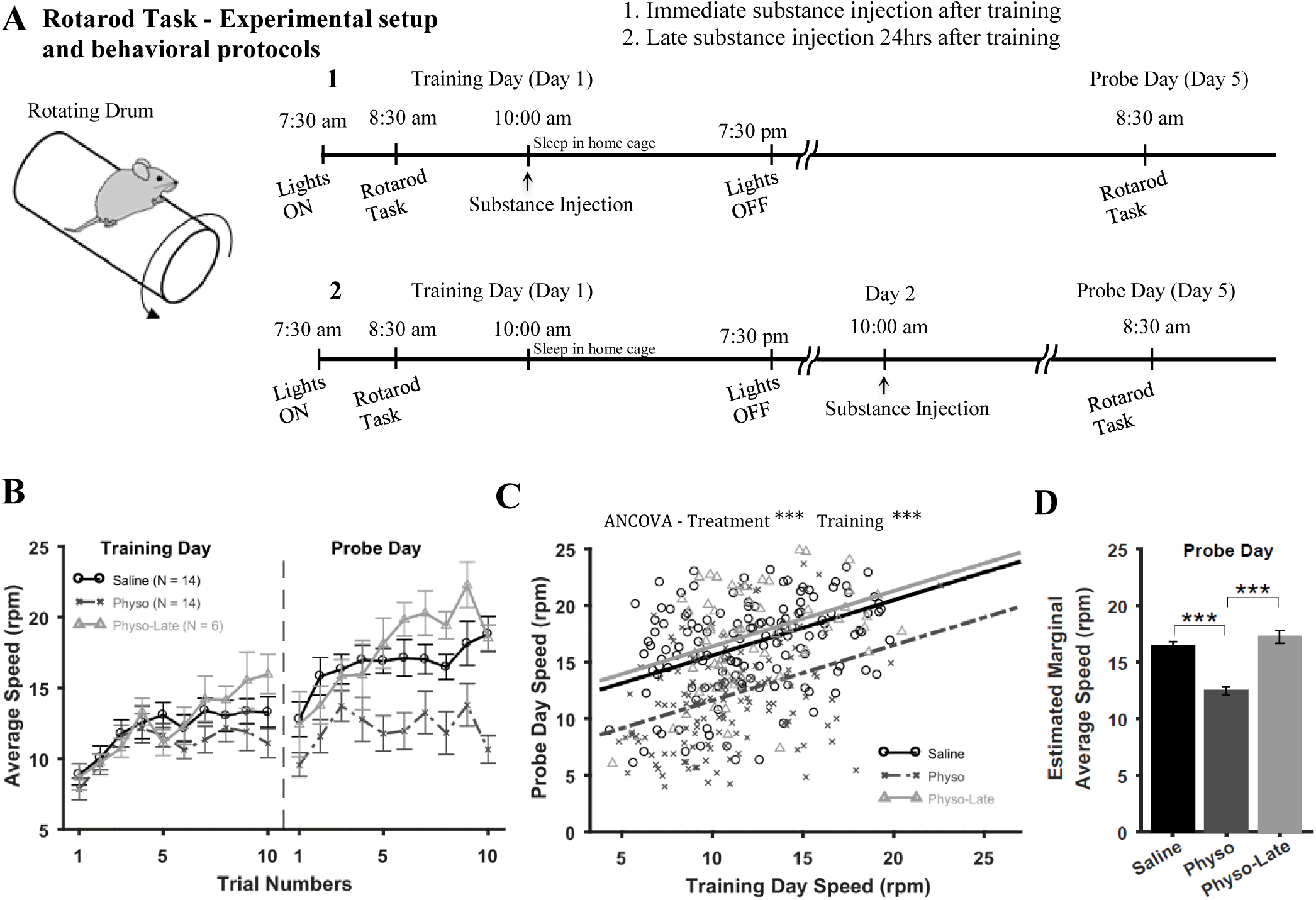
Increasing acetylcholine levels using physostigmine (physo) in early sleep impairs the consolidation of motor memories induced by the rotarod task. **(A)** Experimental setup and behavioral protocol. In the rotarod task, animals learn to run and keep balance on an elevated rotating drum to prevent themselves from falling. Mice were trained on Day 1 (Training day) and retested on Day 5 (Probe day). **(B)** Average speeds (over animals) attained on the rotarod on trials for Training and Probe days. N indicates number of animals in each group. **(C)** ANCOVA results. Scatter plot of Probe Day vs Training Day speeds for each group and fitted lines according to Eq. (1). **(D)** Estimated marginal average speeds on Probe Day for the three groups. Error bars represent SEM.

We used the modified version of the rotarod as described by Shiotuski et al (Shiotsuki H et al. 2010), that emphasizes the learning aspect of the test. A four lane rotarod with automatic timers and fall sensors (Med Associates Inc.) within a test chamber (74 cm / 84 cm / 50 cm) was used. The diameter of drum was 7cm and it was covered with Duct tape to prevent mouse from gripping the surface. Animals were placed on the drum 3 mins before the start of each session for habituation. An accelerating rotarod design was used, in which the rotation speed gradually increased from 4-40 rpm over the course of 5 min. The time latency and the rotation speed were automatically recorded by the photo sensors until the animal was unable to keep up with the increasing speed and fell. Training and testing took place on separate days:

1. *Training Day*. The rotarod training on day 1 consisted of one session (1-1.5h) with 10 trials and an inter trial interval of 3-5 mins. Performance was measured as the average speed that animals achieved during the training session.
2. *Probe Day*. To evaluate long-term memory, 10 trials were again given 5 days after the initial session.

The handling of animals started at 8:00am i.e. moving them from their home cages to the testing room and the rotarod task was started at 8:30am. Thus, all sessions were performed at the same time 8:00-10:00 am in the morning before the animals were returned to home cages to allow them to sleep. Although, the time of training is a non-preferred circadian time, it was chosen for logistic reasons as is described in previous studies (Yang G *et al*. 2014; Nagai H et al. 2017).

#### Skilled-Forelimb Reach Task

The skilled forelimb reaching task (Whishaw IQ 2000) assesses fine/precise co-ordinated movements of forelimb, arm, hand and digits as they work together to retrieve a food item from a shelf. The animals were placed on food restriction 3-4 days before the beginning of training. Prior to food restriction, mice were weighed each day for three days to obtain an average pre-restriction weight. Thereafter they received daily monitoring during the food restriction period to maintain 85% of the average prerestriction weight. For the task, 21 C57/BL6 mice were divided into Control (5 males, 5 females) and Drug (6 males, 5 females) groups by an independent evaluator. On each day of testing the order of the animals to be tested was randomized by the independent evaluator. Thus, throughout the procedures the experimenter was blind to the identity of the two groups.

The test box was clear plexiglass testing chamber (20 cm long, 9 cm wide, and 20 cm high) with a slit (1 cm wide) located in the center of the front wall. A 3cm wide shelf with two divots located aligned with each side of the slot was mounted 1cm above the floor and outside of the front wall. The divots served as receptacles in which 14mg food pellets were placed. The mice were handled daily and were habituated to the testing chamber for a week. Two days before training, the mice were given 10 pellets inside the chamber and then 10 pellets very close to the slit on shelf before the pellets were placed into the divots. The training/testing began on the day the mice were presented with food pellets close to the slit on the shelf. Gradually, food pellets were placed away from the slit and eventually placed in the divots located on the shelf. The training/testing consisted of 10 days with one session each day consisting of 20 trials. A **“**Reach” was scored each time an animal extended its forelimb through the slot. A “Success” was scored if the animal grasped the food, retracted the paw and brought the pellet back to its mouth and consumed it. On each day, immediately after training/testing and before sleep, all mice received either physostigmine or saline and were allowed to sleep. On 9^th^ day of training/testing all mice were filmed with a Panasonic HDC-SDT750 camera at 60 frames per second with an exposure rate of 1ms. Illumination for filming was obtained from a two-arm cold light source (Nikon Inc.). Quality of reaches was assessed from the videos using a standard movement scoring scale (Whishaw IQ 2000). Components of reach that were scored were: hind feet position, forefeet position, sniff, lift, elbow in, advance, pronation, grasp, supination I, supination II, release, and replace. Each movement was scored on a three-point scale; 0, 0.5, and 1; scores representing good, impaired, and absent components respectively. Thus, a high score indicated inferior performance.

### Drugs and Solutions

The drugs Physostigmine Salicylate, Scopolamine Hydrobromide, Mecamylamine Hydrochloride, Oxotremorine M, and Nicotine hydrogen tartrate salt administered intraperitoneally were used in the study. The drugs were obtained from Sigma-Aldrich and dissolved in saline for administration. Saline was injected as a control treatment. Physostigmine was first dissolved in 100% ethyl alcohol to make a stock solution of 10mg/ml and later diluted 1000 times in saline to obtain a final concentration of 0.01mg/ml. All Drugs were prepared just before use or diluted from a stock solution. For the rotarod task, mice were assigned to six groups. Each group received one of the drugs; physostigmine (Physo group), Oxotremorine (Oxo group), Nicotine (Nico Group), Scopolamine and Mecamylamine (Sco+Mec group), and Saline (Saline group). For the skilled reaching task, mice were assigned to two groups with one receiving physostigmine and in the other saline injected after training.

#### Pharmacological Treatment (rotarod task)

For these experiments, five groups were used and the procedure of administering a Training Day and a Probe Day five days later was used (see above):

*(1) Physostigmine group (Physo group).* To increase cholinergic tone during NREM sleep, 14 mice (6 males, 8 females) were administered physostigmine (an acetylcholine esterase inhibitor which indirectly stimulates both muscarinic and nicotinic receptors) intraperitoneally immediately after the first rotarod training session (Gais S and J Born 2004; Jafari-Sabet M et al. 2016; Dhingra D and K Soni 2018). Because it was a single dose study, an acute dose of 0.1mg/kg physostigmine was used which is about ten times the dosage used in a human study (Gais S and J Born 2004). After the training session and drug administration, the animals were transferred to their home cages, where they could sleep. The elimination half-life of physostigmine is documented to be 1.5-2 hours in humans (Hartvig P et al. 1986) and less than 1 hour in rodents (Somani SM 1989), a dose that is enough to stay in the system for the course of early sleep. No obvious drug side effects were observed. For the control condition, 14 mice received saline (S group).
*(2) Physostigmine-late Group (Physo-late 24hrs)*. In order to investigate the role of first sleep session in motor memory consolidation and determine whether or not the performance of the animals on probe day was a result of enduring side-effects from the Day 1 physostigmine treatment, 6 mice (all males) were trained on the rotarod on Day 1 and injected with same dose of physostigmine 24 hours later just before testing.
*(3) Scopolamine/mecamylamine group (Sco+Meca group).* To block cholinergic transmission, we simultaneously administered the muscarinic receptor antagonist scopolamine (0.4mg/kg) and the nicotinic receptor antagonist mecamylamine (3mg/kg) intraperitoneally to 8 mice (4 males, 4 females). We chose relatively low doses of the drugs to avoid strong side effects of cholinergic blockade and to ensure that the substances had largely washed out at the time of probe testing. The half-life in plasma has been estimated at 4.5 ± 1.7 hour for scopolamine (Putcha L et al. 1989) and 10.1 ± 2 hour for mecamylamine (Young JM et al. 2001). Previous studies have shown that these drugs, if not given in combination, do not affect declarative memory encoding or consolidation (Rasch BH et al. 2006). The drugs were given after the end of the rotarod session on Day 1.
*(4) Oxotremorine and nicotine Groups (Oxo and Nico respectively)*. To observe ACh receptor specificity for mediation of motor memory consolidation, we administered either oxotremorine (0.01mg/kg) or Nicotine (2mg/kg), a selective muscarinic and nicotinic ACh receptor agonist respectively, to two separate groups of mice (oxotremorine n=8, 5 males, 3 females, nicotine n=8, 6 males, 2 females) after the rotarod task. The doses were chosen based on studies that have examined the effect of nicotine and oxotremorine on rodents (Power AE et al. 2003). The half-life has been reported as 100 minutes for oxotremorine and 1.5-2 hours for nicotine (Moyer TP et al. 2002; Marchand M et al. 2017). The drugs were administered at the end of training on Training Day.
*(5) Transgenic Groups*. Seventeen transgenic mice, 9 (4 males, 5 females) in the CNO group (Tg-CNO) and 8 (2 males, 6 females) in the saline group (Tg-Saline), were used. The CNO, 5mg/kg (MacLaren DA et al. 2016), was obtained from the Thermo-Fisher Scientific and was dissolved in 10% DMSO then diluted to a final concentration of 0.5 mg/ml CNO with saline. As the onset of the action of CNO begins 1-hour post administration, CNO was injected just before the rotarod training session so that the effect of drug would be manifest in post-training sleep (McCarthy EA et al. 2017; Saund J et al. 2017; Jendryka M et al. 2019).

### Motion quantification with filming in home cage

In a separate cohort of mice (n = 10), sleep recording was done by filming the animals in their home cages after the rotarod training session in order to observe their motion for quantification of post-learning activity. Five mice were given saline and 5 received physostigmine. A Pi camera connected to a Rasp-berry pi computer was attached to the lid of the home cage and 2.5 hours long videos were recorded post training (Singh S et al. 2018). Motion quantification was performed in Matlab^®^ (Mathworks, Natick, MA) by first identifying differences in consecutive frames and then calculating the mean difference over pixels, to obtain an approximation of the overall change in the values of pixels from one frame to the next. The accuracy of motion estimation was verified by manual observation.

### Sleep quantification using electrophysiolog*y*

Four adult C57BL6J mice and one Chat-Cre::CAG-hM4Di mouse were anesthetized with isoflurane (2.5% induction, 1-1.5% maintenance) and implanted with cortical, hippocampal, and muscular electrodes using an aseptic technique. For cortical and hippocampal recording of local field potentials (EEG), bipolar (tip separation = 0.6 mm) and monopolar electrodes made from Teflon-coated stainless-steel wire (bare diameter 50.8 µm) were implanted in the neocortex and in the CA1 hippocampal region using the following coordinates in mm: Secondary motor cortex (M2), AP: 1.7, ML: 0.6, DV:1.1 mm, Lip cortical sensory area (LP), AP: 0.75, ML: 3.0, DV: 1.6, retrosplenial cortex (RS), AP: −2.5, ML: 0.6, DV: 1.1, barrel cortex (BC), AP: −0.1, ML: 3.0, DV: 1.4, and hippocampus (HPC), AP: −2.5, ML: 2.0, DV:1.1 mm. For EMG recording, a multistranded Teflon-coated stainless-steel wire (gauge 40) was implanted into the neck musculature using a 25 gauge needle. The reference and ground screws were placed on the skull over the cerebellum. The other pole of the electrode wires were clamped between two receptacle connectors (Mill-Max Mfg. Corp.) and the headpiece was secured to the skull using metabond and dental cement.

Animals recovered for 10 days after surgery and were then habituated for 5 to 7 days to the recording setup. On baseline days, animals were injected with saline at 8:25 am and moved to the recording setup where their sleep activity was recorded from 8:30 am until 12:30 pm, via a motorized commutator (NeuroTek Inc.). This 4 hr period of baseline recording was repeated for three days. On fourth day, the same steps and the same recording period was used except the animals were injected with physostigmine (dosage 0.1 mg/kg). Local filed potentials and EMG activty were amplified, filtered (0.1-4000 Hz) and digitized at 16 kHz using a Digital Lynx SX Electrophysiology System (Neuralynx, Inc.). The data were recorded and stored on a local PC using Cheetah software (Neuralynx, Inc.). In addition to the electrophysiological recordings, a Pi camera was used to record animal’s behaviour during each session.

#### Data Analysis

The data analyses were performed offline. Sleep scoring was performed in 6-sec long epochs. The raw EMG activity was filtered between 90 Hz and 1000 Hz and then rectified and integrated using 4-sec moving windows. This signal was thresholded to detect periods of immobility. Slow wave power (0.5 to 4 Hz) of the cortical LFP was calculated in each epoch using taper spectral analysis and thresholded using values between 0.04 and 0.1 mV^2^/Hz (depending on the animal) to detect SWA in cortical recording. When the animal was immobile and cortical LFP showed SWA, the epoch was scored as NREM sleep. For detecting REM sleep, the ratio of theta power (6 to 9 Hz) to the total power of hippocampal LFP was calculated in each epoch using taper spectral analysis and when this ratio was above 0.4-0.6 (different for individual animals) and the EMG showed immobility, the epoch was scored as REM sleep. All other epochs were considered as a waking state. The waking and NREM/REM scores were further confirmed using video recording. To investigate the effect of physostigmine on sleep, the duration of each behavioral state in each recording session was normalized to the recording duration to identify differences between physostigmine and saline. NREM sleep latency was calculated as the time between the onset of recording and the onset of first NREM sleep epoch longer than 20 seconds. Moreover, slow wave power (0.5 to 4 Hz) of cortical signal was averaged across all NREM sleep epochs in the first hour of recording and then compared between physostigmine and saline. For saline injection, sleep structure, NREM sleep latency and SW power were averaged across three baseline days for each animal. Power in other frequency bands (theta, alpha, beta, and slow and fast gamma) was also determined and presented in supplementary figures.

For detecting spindle events, following Philips et. al., the M2 cortical signal was filtered between 8 and 18 Hz and then rectified (Phillips K et al. 2012). The envelope of the rectified signal was then calculated using cubic interpolation of its maxima. Putative spindle events were detected where the envelope signal was higher than 3.5 standard deviations (SD) from its mean. The onset and offset of spindle events was determined where the envelope signal was higher than 2.5 SD from its mean. Spindles shorter than 500msec or longer than 2sec were rejected. Spindles that occurred within 100 msec of each other were combined to be considered a single event.

### Sleep deprivation

To test the effect of sleep deprivation on motor memory consolidation, we used 16 mice which were randomly assigned to Control (n = 6, 2 males, 4 females) and Sleep-Deprived (n = 10, 5 males, 5 females) groups. Animals in Sleep-Deprived group were kept awake for 1-hour post training on Day1 while ones in the Control group were put in their home cages and allowed to sleep. To keep animals awake, when they showed signs of sleep featuring little motion and closed eyes, they were aroused by gently moving them by hand, rearranging their nest, or lifting and lowering their cage to keep them awake. Animals that were awake were not disturbed.

### Statistical Analysis

Statistical analysis was done using Matlab R2016b. Data are presented as mean ± the standard error of mean (S.E.M). The following statistical tests were used; paired and two-sample Student’s t-test, Kolmogorov-Smirnov test, analysis of covariance (ANCOVA), and analysis of variance (ANOVA) with repeated measures. An alpha value of 0.05 was used to determine significance.

For rotarod experiments, since we measured rotarod performance before and after substance administration (experimental design of pretest-posttest type), we used a one-way ANCOVA to statistically determine the effect of Treatment (drug administration) on Probe Day speed by using Training Day speed as a covariate (Huck SW and RA McLean 1975). We used the following fixed-effects linear model for ANCOVA;

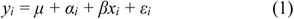

where *y* is the response variable i.e., rotarod speed on probe day, subscript *i* indicates the treatment group, *x* is training day speed (covariate), *μ* is the main intercept, α is the treatment specific intercept, *β* is the slope, and *ε* is the error term. Note that the slope is same for all groups i.e., assumption of parallel fitted lines. Initially, we included the interaction term “*α_i_x_i_*” in our model (to account for different slopes of separate fitted lines) but determined using Matlab’s “step” function on linear regression models that this term was unnecessary for predicting probe day speeds (for all ANCOVA tests). The step function sequentially adds and eliminates terms in the model to reduce terms while preserving predictive accuracy. In our model, we also did not include a term for Sex effects because we had unequal number of males and females in individual groups. Furthermore, from a preliminary analysis of rotarod performance for 74 mice (39 males and 35 females) which received identical training on Training Day, we found no differences in performance between males and females. We performed a repeated measures ANOVA with Sex as “between-subjects” factor and Trial as “within-subjects” factor and found no significant effect of Sex (F_1,72_ = 0.02, p = 0.877) or significant Sex by Trial interaction (F_9,648_ = 1.19, p = 0.307).

## RESULTS

### Physostigmine injection after a motor learning task impairs the consolidation of motor memories

Mice successfully completed the rotarod and skilled forelimb reaching tasks, which feature, rapid learning (10-20 trials within a single training session), allowing assessment of the effects of manipulations during relatively immediate post-training sleep, (Yang G *et al*. 2014) and more gradual learning (multiple training sessions each with 20 trials spread over 2-6 days), with potentially multiple cycles of encoding and post-training sleep, respectively. The following results describe the effect of post-training and pre-sleep injections of physostigmine, an acetylcholinesterase inhibitor that prevents the breakdown of ACh, thus increasing cholinergic tone (Aquilonius SM and P Hartvig 1986; Pubchem 2018).

#### Rotarod Task Results

In the rotarod task (Fig. 1A), mice displayed improved balance on the elevated rotating drum as witnessed by the reduced probability of their falling. The maximum speed they attain just before a fall was used as a measure of learning and subsequent retest performance. On day 1 (Training Day) mice were given 10 trials of rotarod experience early in the morning (8:30am – 10:00am). Immediately after training, one group received saline and the other physostigmine and they were returned to their home cages. Mice in a third group (Physo-Late) were also returned to their home cages but physostigmine was injected 24 hours after the training. On the Training Day, all groups displayed improved performance across trials by remaining on the rotarod for progressively longer periods of time i.e., at higher speeds (Fig. 1B). To assess whether the mice retained motor performance acquired on the Training Day, they again received 10 trials of rotarod training on day 5 (Probe Day), at the same time of the light/dark cycle on which initial training had occurred (Fig. 1A). The Saline and Physostigmine-Late treatment groups displayed continued improvement in performance across trials but the Physostigmine group displayed performance that appeared to be no better than that they had displayed on the Training Day.

These results were confirmed by ANCOVA using the fixed-effects linear model of Eq. (1) – Fig. 1C. There was a significant effect of Treatment (F_2,336_ = 38.77, p < 0.0001) as well as Training (F_1,336_ = 57.88, p < 0.0001). Post hoc multiple comparisons with Bonferroni correction revealed a significant difference in population marginal means of Saline and Physo-Late groups from that of Physostigmine group (p < 0.0001 for both comparisons, Fig. 1D). Saline and Physo-Late groups were not different from each other (p = 0.691). These results thus suggest that the Saline and Physostigmine-Late groups displayed enhanced motor memory for the task relative to the Physostigmine group and that the immediate post-training sleep is important for performance retention.

#### Skilled Forelimb Reach Task Results

In the single pellet reaching task (Fig. 2A) animals coordinated their arm and hand movements to reach through a slot to obtain a food pellet for 10 days, and on each day received either saline or physostigmine immediately after training (Fig. 2A). Mice injected with physostigmine were slower in learning the task, as witnessed by a smaller number of animals learning the task within the first 5 days compared to those injected with saline (Fig. 2B, an association between substance injected and success was observed on days 3 and 4 with Chi square independence test, χ^2^(1) = 10.8315, p=0.00099 and χ^2^(1) = 4.492 p=0.0341 respectively). A successful animal was defined as one that completed 20 trials on that particular day with at least one successful reach. Next, we compared the percentages of successful reaches (Fig. 2C) using repeated measures ANOVA with Treatment (Saline or Physostigmine) and Sex as “between-subjects” factors and Day as “within-subjects” factor. There was no effect of Treatment (F_1,17_ = 3.10, p = 0.096) and Sex (F_1,17_ = 1.56, p = 0.235) but that of Day was significant (F_8,136_ = 28.83, p < 0.0001). There was also significant Treatment by Day interaction (F_8,136_ = 2.51, p = 0.014) but Treatment by Sex (F_1,17_ = 0.05, p = 0.822), Sex by Day (F_8,136_ = 0.88, p = 0.539) and Treatment by Sex by Day (F_8,136_ = 0.47, p = 0.878) interactions were not significant. Post hoc comparisons (Treatment-by-Day) with Bonferroni correction revealed significantly smaller percentage of successful reaches for the physostigmine group for days 7 and 8 (p=0.01 and p=0.02 respectively). These results show that the administration of physostigmine immediately after reach training retarded task acquisition and performance regardless of sex.

**Figure 2.**
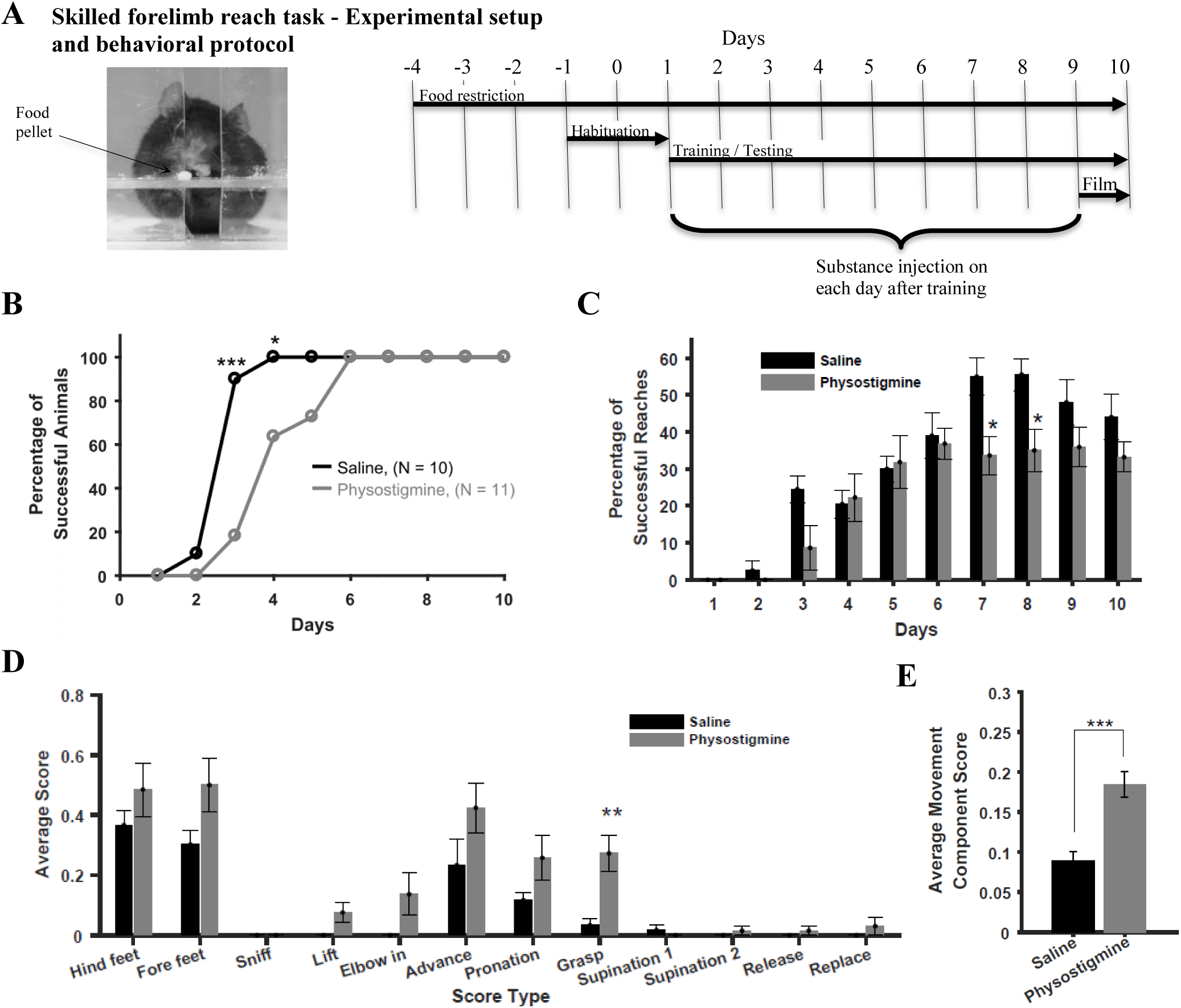
Increasing acetylcholine levels using physostigmine in early sleep impairs the consolidation of motor memories in the skilled forelimb reach task. **(A)** Experimental setup and behavioral protocol. Animals learn to reach through a slot and grasp a food pellet placed on a platform. Mice were trained/tested for 10 days. They were filmed on 9^th^ day and reaches were scored from the videos. **(B)** Percentage of successful animal vs days. A “successful” animal completed 20 trials with at least one successful reach and grasp. **(C)** Percent single reach successes (averaged over animals) vs days. **(D)** Quality of reaches was assessed from video recordings done on day 9 for the 12 component movements of a reaching action. A higher score indicates inferior performance. **(E)** Average component scores (over animals) for the physostigmine group were significantly higher than those for the saline group. All Data are presented as Mean±SEM.

To assess the effect of physostigmine injections on the quality of reaches, the video recordings on day 9, from the first 3 reaches, 12 components of movements were scored on a 0, 0.5, and 1 scale (good, impaired, and absent performance). A higher score thus indicates inferior performance (Whishaw IQ et al. 2008; Whishaw IQ et al. 2017). To compare scores of the two groups, we performed repeated measures ANOVA with Treatment (Saline or Physostigmine) and Sex as “between-subjects” factors and Movement Component as “within-subjects” factor. The effect of Treatment was significant (F_1,17_ = 10.34, p = 0.005) as well as that of Movement Component (F_11,187_ = 24.07, p < 0.0001) but that of Sex was not significant (F_1,17_ = 0.82, p = 0.377). There was no significant Treatment by Component (F_11,187_ = 1.65, p = 0.087), Treatment by Sex (F_1,17_ = 1.65, p = 0.217), Sex by Component (F_11,187_ = 0.38, p = 0.962), and Treatment by Sex by Component (F_11,187_ = 1.04, p = 0.409) interactions. Post hoc comparisons (Treatment-by-Component) with Bonferroni correction revealed a higher mean score for the grasp component for the Physostigmine group relative to the Saline group (p = 0.004). In addition, average component scores (over components and animals) were higher for the Physostigmine group than for the Saline group revealed with a two-sample Student’s t-test (t_754_ = −4.711, p < 0.0001, Fig. 1E). These results thus show that as mice in the physostigmine group acquired the task, their motor performance was impaired compared to the saline group on both end point measures of performance and the movement elements of performance.

### Physostigmine injection after rotarod training altered sleep structure

To investigate how physostigmine injection post-training impaired subsequent motor performance, we asked whether physostigmine injection altered post-training activity/sleep by injecting separate cohorts of mice with saline or physostigmine after rotarod training, before being returned to their home cages. The activity/sleep analysis via video monitoring provided a motion measure (Fig. 3A) as quantified by the number of pixels that changed between consecutive video frames (Singh S *et al*. 2018). The record showed that mice in the physostigmine group had increased activity levels and showed delayed onset of rest/sleep. For each mouse, the cumulative history of motion/rest periods was determined. The average cumulative duration of motion for mice in the saline group was significantly smaller and their immobility periods were longer compared to mice in the physostigmine group (two-sample Kolmogorov-Smirnov test, p < 0.0001, Fig. 3B). The physostigmine group’s relative higher activity and shorter episodes of rest suggests that physostigmine injections degraded sleep in the mice.

**Figure 3.**
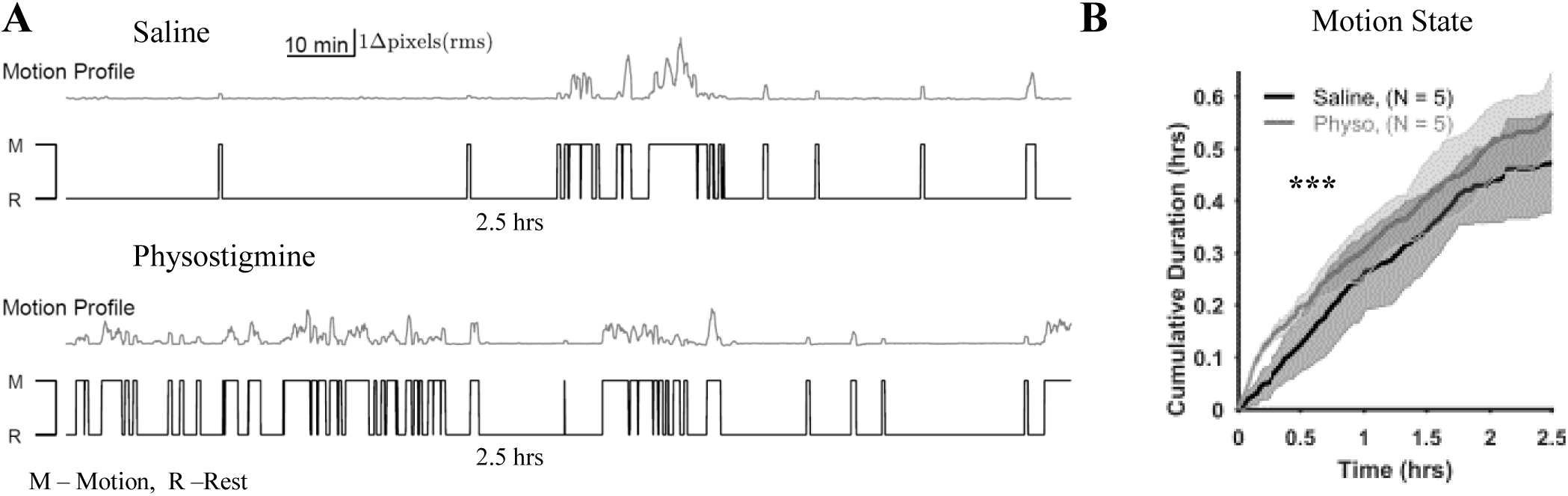
Physostigmine altered sleep structure. Quantification of motion in home cage using video monitoring after rotarod training. **(A)** Representative raw traces of rms value of change in number of pixels in video frames versus time for mice injected with saline and physostigmine (physo). Mean of the whole trace was used as a threshold to classify motion and rest periods shown in black traces below each raw trace. **(B)** Average cumulative motion versus time. For each mouse, the cumulative sum was determined from motion/rest traces. Shaded regions show SEM. The mean curve for Saline group is significantly right shifted compared to Physo group (two-sample Kolmogrov-Smirnov test, p < 0.0001) suggesting reduced motion of mice in Saline group relative to the Physostigmine group.

### Physostigmine delays NREM sleep onset and reduces slow wave sleep power

The examination of how physostigmine alters sleep structure used electrophysiology to monitor EEG in one group of mice (N = 5). We implanted electrodes in the trapezius muscle (in the neck), hippocampus (HPC), secondary motor cortex (M2), cortical lip area (LP), and the retrosplenial and barrel cortices (RS and BC). Local field potentials (LFPs) were recorded after injecting saline and on a subsequent day after injecting physostigmine (Fig. 4A). Mice were injected at 8:25am and signals were analysed for four hours post-injection from 8:30am. In both hippocampal and cortical spectrograms, we observed visible reduction in power spectral density (PSD) in the slow wave band within the first hour after physostigmine injections (Fig. 4A). For further comparisons, we first determined sleep structure by finding durations of wakefulness, REM sleep, and NREM sleep using EMG signals and spectrogram of hippocampal and cortical (from M2) LFPs (Fig. 4A and 4B). Compared to saline injections, sleep structure after physostigmine injections was altered in the first hour of recording. After the first hour, cumulative duration curves for saline and physostigmine injections were the same for awake, NREM sleep, and REM sleep conditions (Fig. 4B). After physostigmine injections mice had delayed onset of NREM sleep (Fig. 4A and 4B), time spent awake was increased (paired Student’s t-test, t_4_ = −4.14, p = 0.014, Fig. 4C), and the NREM sleep period was shorter (paired Student’s t-test, t_4_ = 4.32, p = 0.012, Fig. 4C). REM sleep was minimal during the first hour and not different for physostigmine and saline injections. The percentage of time spent in awake, NREM and REM sleep conditions was not different during the 2^nd^, 3^rd^ and 4^th^ hours (supplementary Fig. S2G). NREM sleep latency was longer after physostigmine (paired Student’s t-test, t_4_ = −3.52, p = 0.024, 31.17±3.26 mins for physo vs 16.70±2.76 mins for saline, Fig. 4D) and slow-wave power within the first hour after the onset of NREM sleep was reduced in the hippocampus (paired Student’s t-test, t_4_ = 6.42, p = 0.003, Fig. 4E) and cortex, (paired Student’s t-test, M2 - t_3_ = 11.74, p = 0.0013, LP- t_2_ = 12.86, p = 0.006, RS-t_2_ = 14.09, p = 0.005, and BC-t_3_ = 6.23, p = 0.008, Fig. 4E). Theta (4-9Hz), alpha(9-15Hz), and beta (15-30Hz) powers in the hippocampus were also reduced after physostigmine injections within the first hour after the onset of NREM sleep but changes were variable in the cortices (supplementary Fig. S1 A-E) i.e., there was reduction in power in alpha, beta, and slow gamma bands for M2, no changes in power for LP, reduction in power in alpha and beta bands for RS, and reduction in power in beta band for BC. Changes in power with physostigmine injections in all frequency bands were also variable in the 2^nd^ and 3^rd^ hours after the onset of NREM sleep. Hippocampal alpha and beta band power reduced with no effect on slow wave power and power in other bands while power was reduced in some frequency bands in cortical areas M2, RS, and LP with no change for BC (supplementary Fig. S2). The onset of thalamocortical spindles in M2 was delayed after physostigmine injections and their number was significantly reduced (Fig. 4F). These results suggest that physostigmine injection delayed the onset of NREM sleep and reduced its quality (lower power in slow wave and higher frequency bands) in the first recording hour.

**Figure 4.**
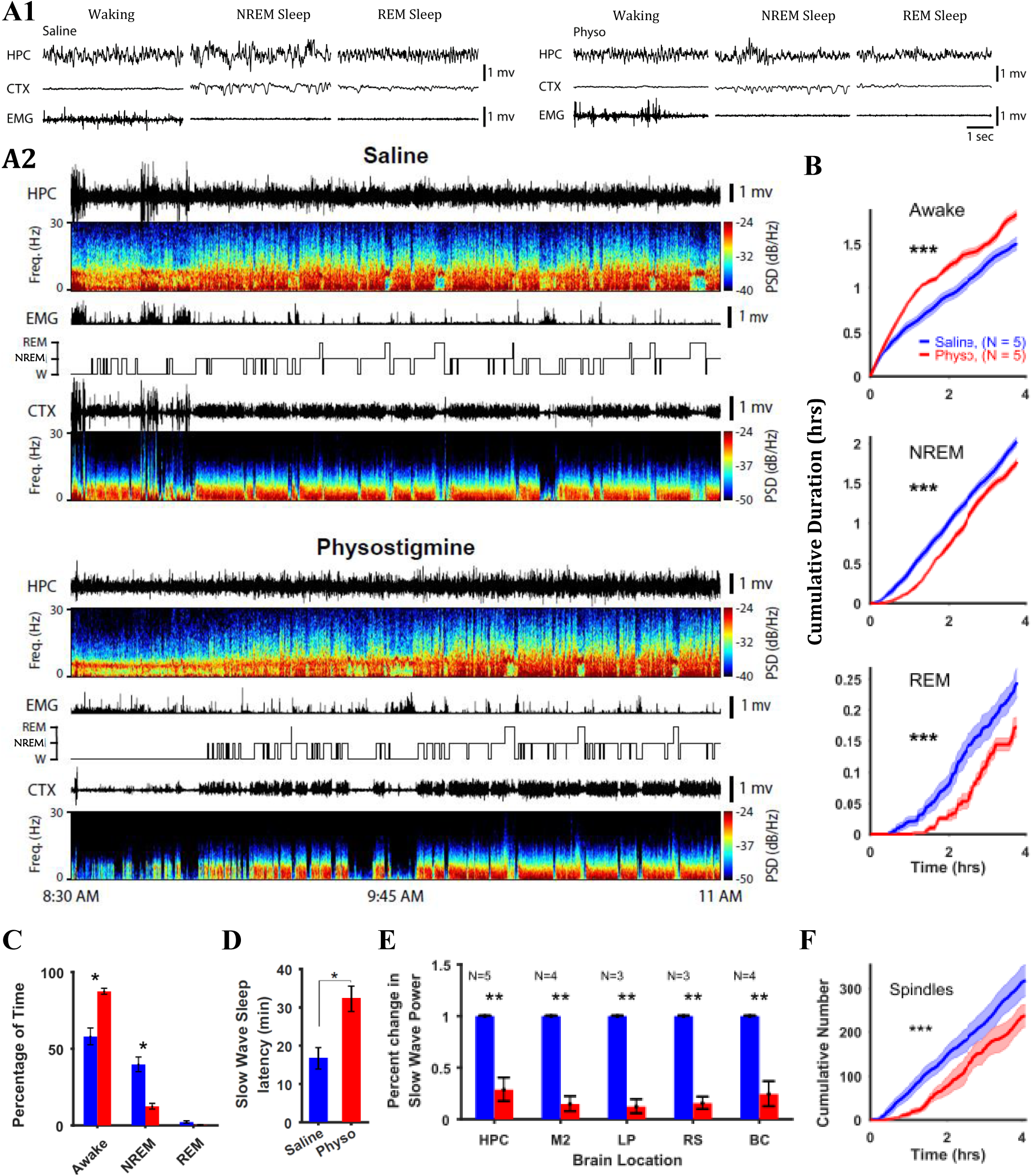
Increasing acetylcholine levels via physostigmine (physo) reduces NREM sleep and slow-wave power in the first hour of sleep post injection. **(A)** Representative raw local field potentials and EMG during waking and NREM sleep after saline (left) and physostigmine (right) injections (A1). Representative hippocampal (HPC) and cortical (CTX – from M2, secondary motor area) local field potentials (LFP), electromyography (EMG) signal, hypnogram, and spectrogram (A2). In the hypnogram, REM, NREM, and W represent states of animals i.e. REM sleep, NREM sleep, and wake periods respectively, which were scored from EMG traces and spectrogram of hippocampal and cortical LFPs. PSD is power spectral density. Note that the hypnogram for physostigmine shows more fragmented sleep structure compared to that for saline. **(B)** Cumulative duration vs time of recording for awake, NREM, and REM sleep states. Difference in states between animals injected with physostigmine and saline lies mostly in the first hour post injection. Mean cumulative curves for all states were significantly shifted between animals injected with saline and physostigmine (Kolmogorov-Smirnov test, p < 0.0001 for all comparisons) **(C)** Comparison of sleep structure between saline and physo groups in the first hour after injection. Waking activity in physo group was higher (paired Student’s t-test, t_4_ = −4.14, p = 0.014) while NREM sleep was reduced (paired Student’s t-test, t_4_ = 4.32, p = 0.012). **(D)** NREM sleep latency increased in physo group compared to saline group (paired Student’s t-test, t_4_ = −3.52, p = 0.024). **(E)** Slow wave power during NREM sleep in the first hour of recording is reduced in physo group compared to saline group in HPC and cortical areas (M2 – Secondary motor area, LP – Lip primary sensory area, RS – retrosplenial cortex, and BC – barrel cortex) revealed with paired Student’s t-test. N indicates number of samples (total 5 animals). **(F)** Cumulative number of thalamocortical spindles identified from M2 cortical LFP during the NREM sleep period was smaller in physo group compared to saline. All Data are presented as Mean±SEM.

### One-hour sleep deprivation (delayed sleep) after rotarod training does not impair consolidation of motor memories induced by the rotarod task

Our investigation of whether the observed differences in rotarod performance were due to the sleep disruptive effects of physostigmine vs increasing acetylcholine levels (Figs. 3 and 4), used measures from a separate cohort of animals, in which one group was sleep deprived for the first hour post rotarod training and a Control group was left undisturbed (Fig. 5A). Performance assessment indicated that mice in both groups improved on the Training and Probe days (Fig. 5B). An ANCOVA (Fig. 5C) confirmed no effect of Treatment (F_1,157_ = 0.12. p = 0.725) but a significant effect of Training (F_1,157_ = 11.90. p < 0.0001). The estimated marginal average speeds for the two groups were similar on the probe day (Fig. 5D). We also quantified the motion of animals post-training with home-cage video recording (Fig. 5E) and observed that sleep deprived animals had lower overall motion (two-sample Kolmogrov-Smirnov test, p < 0.0005), suggesting that they slept more compared to controls. These results suggest that one-hour of sleep deprivation post motor training does not impair the consolidation of motor memories i.e., delayed sleep after motor training allows motor memory consolidation.

**Figure 5.**
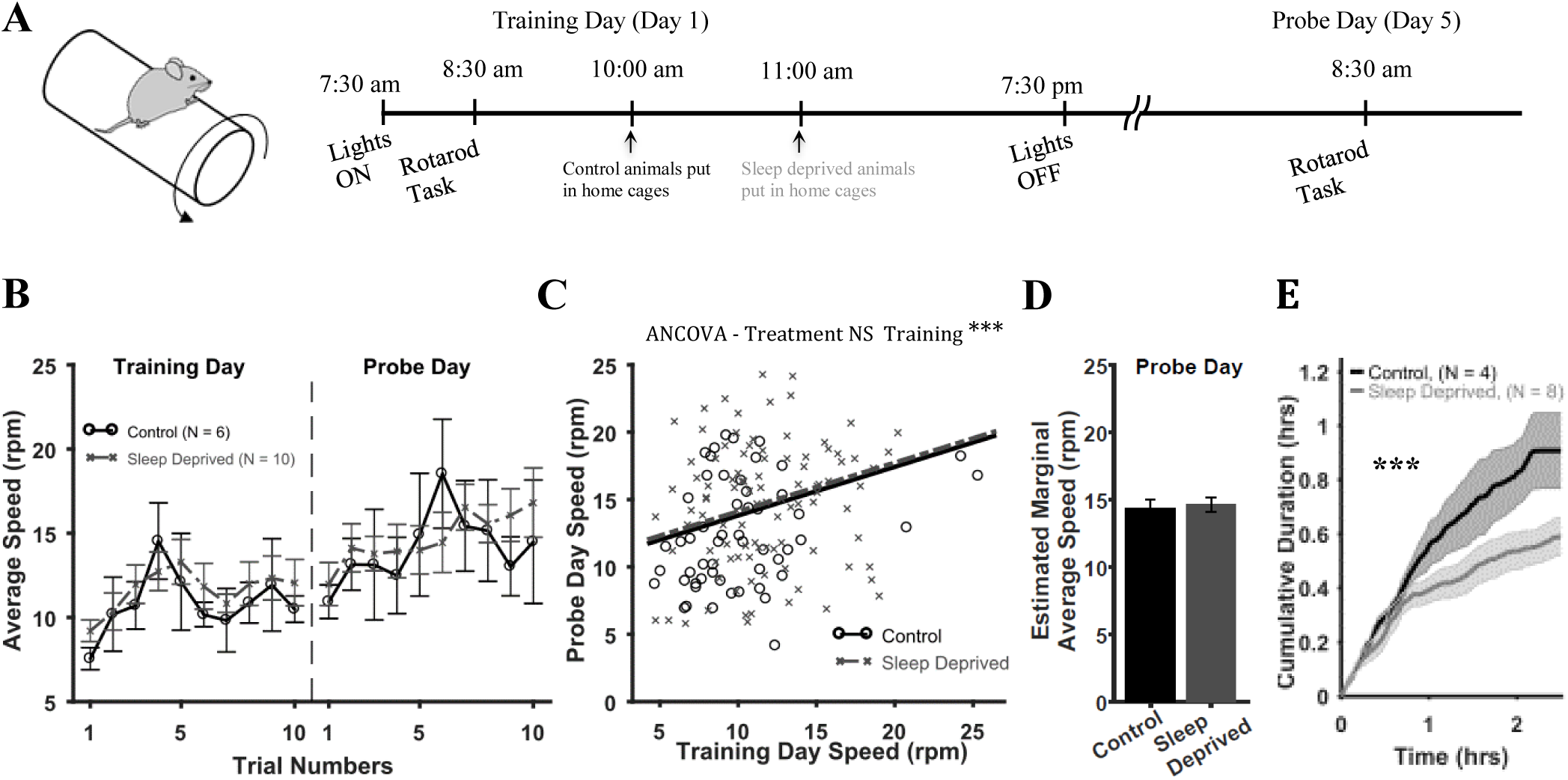
1-hr sleep deprivation post rotarod training does not impair motor memories induced by the rotarod task. **(A)** Control animals were placed in their home cages at 10am while sleep deprived animals were kept awake and put in home cages at 11am **(B)** Average speeds (over animals) attained on the rotarod as a function of trials. N indicates number of animals in each group. **(C)** ANCOVA results. Scatter plot of Probe Day vs Training Day speeds and fitted lines for the two groups according to Eq. (1). **(D)** Estimated marginal average speed on Probe Day was similar for both groups. **(E)** Cumulative motion versus time. Sleep deprived animals had smaller motion. Shaded regions and error bars represent SEM.

### Chemogenetic inactivation of cholinergic neurons during early sleep impairs the consolidation of motor memories

Alternatively to increasing acetylcholine levels using physostigmine, here, we investigated the effect of further reducing acetylcholine levels during early sleep on motor memory consolidation using transgenic Chat-Cre::CAG-hM4Di mice (Sternson & Roth, 2014; Madisen et al., 2010). These mice express inhibitory (hM4Di) DREADDs (Zhu H et al. 2016) within cholinergic neurons, and following the administration of the drug clozapine-*N*-oxide (CNO), presynaptic release of neurotransmitter acetylcholine in cholinergic neurons decreases. Since CNO exerts its maximum effects an hour to an hour- and-half after the administration (McCarthy EA *et al*. 2017; Saund J *et al*. 2017; Jendryka M *et al*. 2019), we injected CNO just before the rotarod training in mice belonging to the Chat-Cre::CAG-hM4Di CNO group (Fig. 6A) to compare them with a Control group of Chat-Cre::CAG-hM4Di mice injected with saline (Fig. 6A). The performance of mice in both groups improved with trials on Training as well as Probe Days (Fig. 6B). However, mice in the Saline group had better performance on Probe Day. With ANCOVA (Fig. 6C) we found a significant effect of Treatment (F_1,167_ = 10.76, p = 0.0013) as well as Training (F_1,167_ = 20.24, p < 0.0001). The Saline group had a significantly higher estimated marginal average speed for Probe Day compared to CNO (Fig. 6D). These results suggest that the action of CNO on transgenic mice during early sleep appreciably affected rotarod performance.

**Figure 6.**
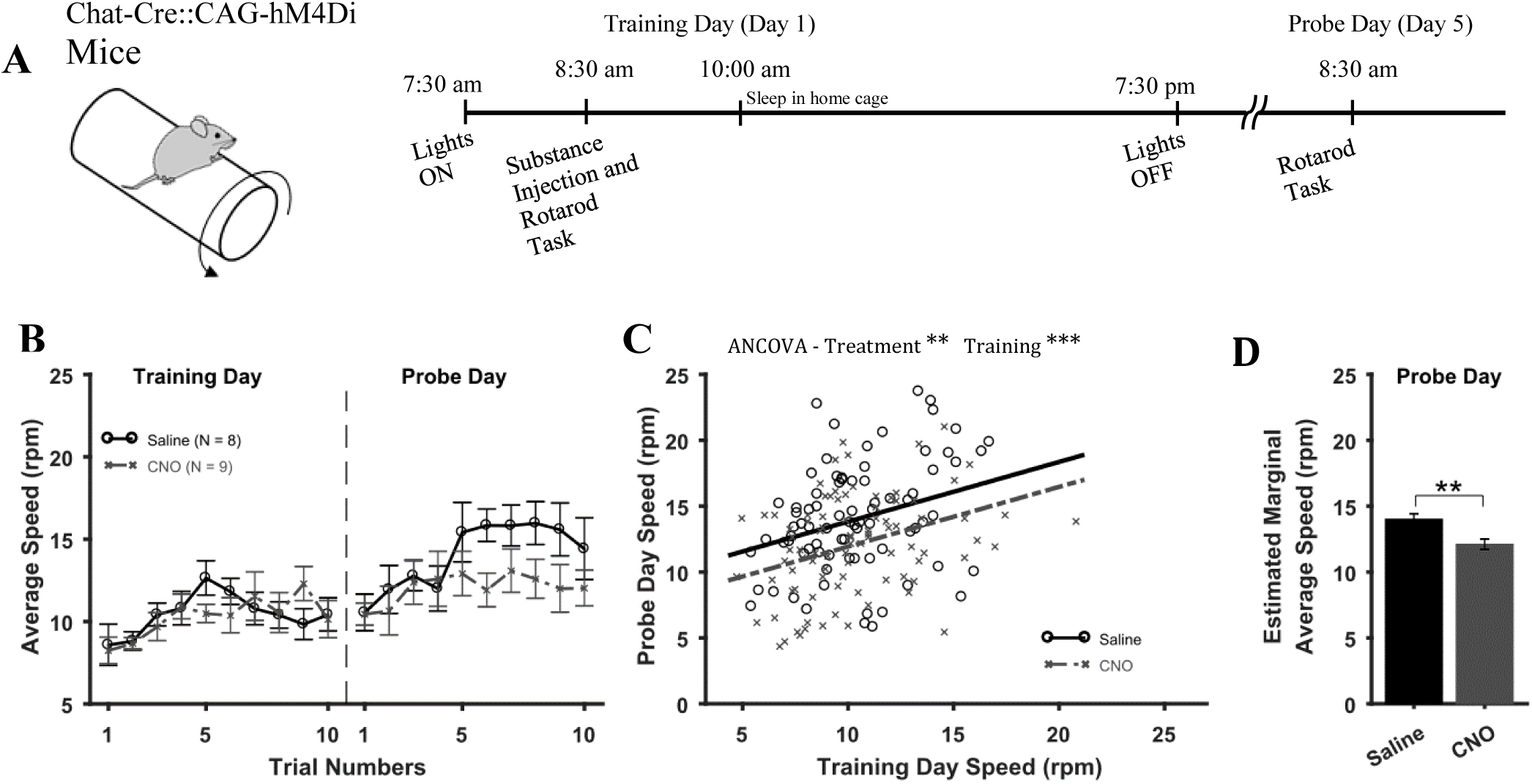
Decreasing acetylcholine levels in early sleep impairs motor memories of the rotarod task. **(A)** Chat-Cre::CAG-hM4Di transgenic animals were injected with either saline or clozapine-*N*-oxide (CNO) before starting rotarod training because CNO exerts its maximum effect an hour to an hour and a half after administration. **(B)** Average speeds (over animals) attained on the rotarod versus trials. N indicates number of animals in each group. **(C)** ANCOVA results. Scatter plot of Probe Day vs Training Day speeds for each group and fitted lines according to Eq. (1). **(D)** Estimated marginal average speeds on Probe Day for the two groups. Error bars represent SEM.

### Activation of muscarinic ACh receptors during early sleep impairs motor memory consolidation

Because post-training physostigmine indirectly activates both muscarinic and nicotinic ACh receptors, we investigated the individual involvement of ACh receptors on motor memory consolidation. Based on the anatomical distribution of ACh receptors in the brain, we hypothesized that, similar to previous studies, the muscarinic ACh receptors would play a larger role in motor memory consolidation than nicotinic ACh receptors (Power AE *et al*. 2003; Brown DA 2010; Hut RA and EA Van der Zee 2011). To activate muscarinic and nicotinic ACh receptors selectively, the respective agonists oxotremorine and nicotine were injected immediately after rotarod training in separate groups of mice (behavioral protocol 1, Fig. 1A). In a third group, Scopolamine and Mecamylamine were injected for combined blockade of muscarinic and nicotinic ACh receptors. These three groups were compared with the Saline group (Fig. 7A). An ANCOVA (Fig. 7B) found a significant effect of Treatment (F_3,375_ = 6.37, p = 0.0003) as well as Training (F_1,375_ = 28.50, p < 0.0001) on Probe Day speeds. Post hoc comparisons with Bonferroni correction showed a significantly higher estimated marginal average speed for Saline group compared with Oxotremorine group (p = 0.0001) but rest of the comparisons yielded no significant difference (Fig. 7C). Oxotremorine injections and activation of muscarinic receptors in the immediate sleep after training impaired rotarod performance on Probe Day.

**Figure 7.**
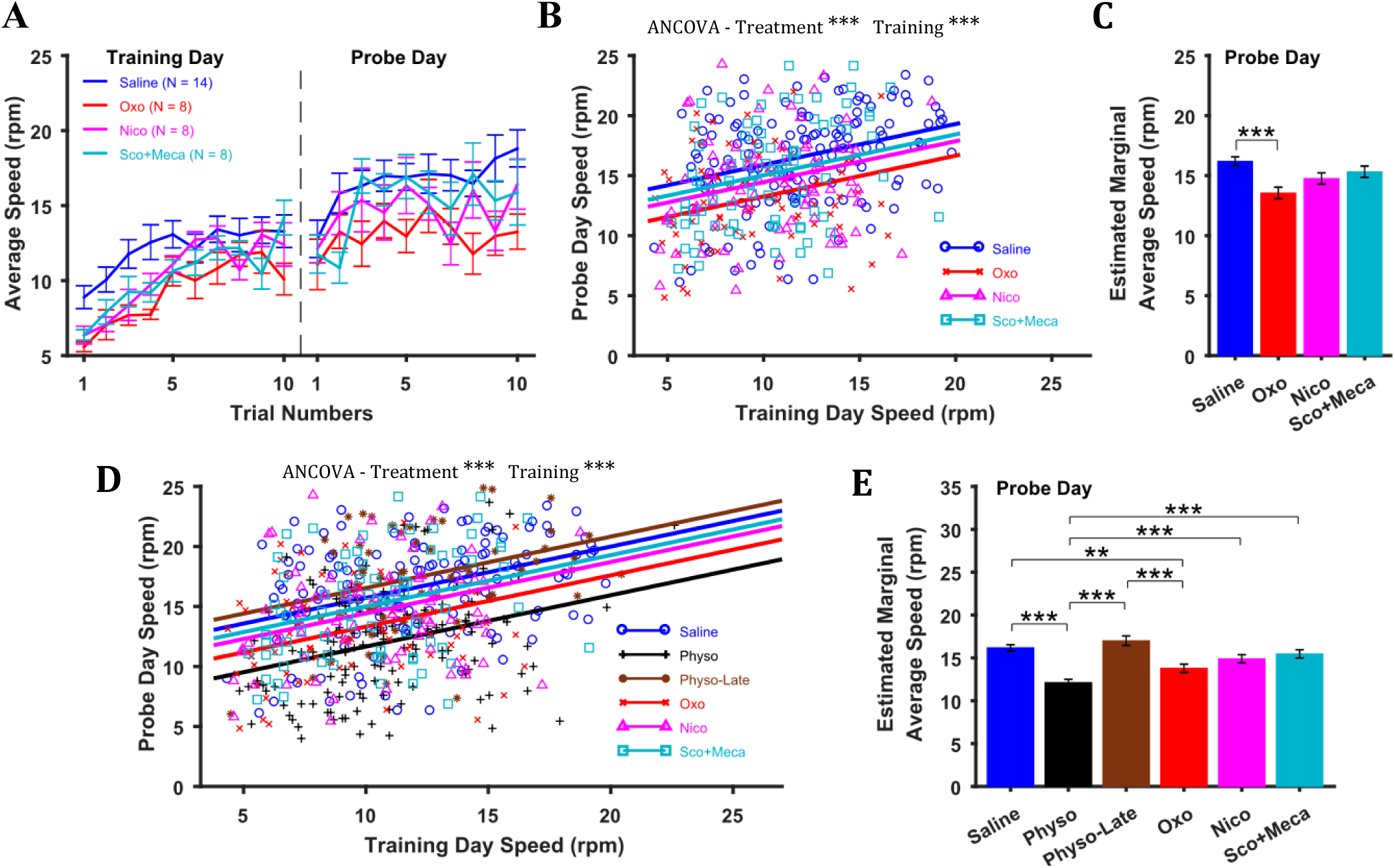
Activation of muscarinic acetylcholine (ACh) receptors during early sleep impairs motor memory consolidation. **(A)** Average speed improvement (over animals) on the rotarod. N indicates number of animals in each group. Saline group is same as in Fig. 1. **(B)** ANCOVA results. Scatter plot of Probe Day vs Training Day speeds and fitted lines for the four groups according to Eq. (1). **(C)** Estimated marginal average speed on Probe Day for the four groups. **(D-E)** Same as (B) and (C) but results of ANCOVA for all groups combined. Error bars represent SEM.

In addition, an ANCOVA on all groups together (Fig. 7D) found a significant effect of Treatment (F_5,573_ = 17.92, p < 0.0001) and Training (F_1,573_ = 69.37, p < 0.0001) on Probe Day speeds. Post hoc comparisons with Bonferroni correction showed that mice in Physo and Oxotremorine groups had lower estimated marginal average speeds compared to all other groups (Fig. 7E). These results corroborate our previous findings that physostigmine and oxotremorine injections affect Probe Day rotarod performance.

## DISCUSSION

We investigated how altering ACh levels during early sleep affects motor memory consolidation as inferred from performance on motor tasks. Supporting previous studies of episodic memory (Hasselmo ME 1999; Gais S and J Born 2004; Hasselmo ME and J McGaughy 2004), we found that low levels of ACh during immediate post-training NREM sleep contributes to motor memory consolidation. Physostigmine given immediately after training to increase ACh levels delayed the onset of NREM sleep, reduced slow-wave power in the cortex and hippocampus, and impaired post-sleep motor performance. Physostigmine given 24 hrs after training as a control procedure, after the first sleep session was over, did not affect motor performance. To rule out that sleep disruption by physostigmine might not be the only cause of impaired motor performance, supporting experiments confirmed that one-hour sleep deprivation (or delayed sleep) after training does not affect motor performance. Furthermore, CNO injection into transgenic mice, which might be expected to decrease acetylcholine levels, affected motor performance. A comparison of the effects of agonists and antagonists of muscarinic and nicotinic ACh receptors showed that relatively selective activation of muscarinic receptors during early sleep impairs motor performance. Taken together, these experiments show that inactivation of muscarinic receptors due to natural lowering of ACh levels during early sleep contributes to the consolidation of motor memory.

It is well known that motor skills show the greatest improvement if intermittent training is given (Karni A et al. 1998; Buitrago MM et al. 2004; Peters AJ et al. 2014). A learning pattern, in which daily improvement is preserved and then enhanced on successive days of training, is likely enabled by a post training period of consolidation. Our results from animals injected with saline after training in both the rotarod and the reach tasks reflect a similar pattern of day to day improvement. For the rotarod task, the improvement obtained on the first day was preserved and augmented on a subsequent probe day. Similarly, for the skilled reach task there was a day to day improvement in performance which eventually plateaued after a number of days of training. The expectation underlying the experiments was that post training sleep contributes to the consolidation that underlies daily improvement in motor performance. Alternatively, for animals injected with physostigmine motor performance was adversely affected. For the rotarod task, performance of Physostigmine group on the probe day was similar to that on the training day with a rapid improvement in the first three trials but followed by a plateau in later trials. For the skilled reach task, the rate of learning was slower, but performance plateaued after day 6. Physostigmine injections therefore disrupted post training procedural memory consolidation.

We obtained support for the idea that it is the lower ACh levels during early sleep that is positively related to enhanced motor performance. First, we used physostigmine to increase ACh levels in post training sleep to find it had negative effect on subsequent performance. We did not measure ACh levels directly but inferred from sleep changes that cholinergic tone was increased. In the first hour of sleep, post injection, NREM sleep onset was delayed and there was a reduction in power in slow-wave and higher frequency bands (in hippocampus and neocortices), consistent with half-life of physostigmine in rodents which is less than an hour (Somani SM 1989). Previous studies have reported similar changes in sleep and frequency patterns following manipulation of cholinergic tone (Sitaram N et al. 1976; Vanderwolf CH et al. 1993; Dringenberg HC and CH Vanderwolf 1996, 1997; Gais S and J Born 2004; Jacobson TK et al. 2013; Anaclet C et al. 2015). We then asked whether the impairment in motor performance resulting from physostigmine injections was due to an increase in acetylcholine levels or solely due to sleep disruption. We ruled out the later possibility by showing that one hour of sleep deprivation i.e. delayed sleep, post training does not impair motor performance. Our results agree with Nagai et al (2017) who showed that 7 hours of sleep deprivation does not affect rotarod performance 24 hrs later (Nagai H *et al*. 2017). In contrast, a study by Yang et al (2014) reported that when animals were sleep deprived for 7 hours after training, rotarod performance is reduced 24 hrs and 5 days later (Yang G *et al*. 2014). Nagai et al attributed the discrepancy between their results with Yang et al to differences in laboratory environments. The discrepancy between our results and those of Yang et al might also be due to differences in the duration of sleep deprivation and the rotarod apparatus used (large vs small diameter drums). In our study, one hour of sleep deprivation might not have been enough to see adverse effects on motor memory consolidation. Nevertheless, given the results of Nagai et al, we do not attribute the effects of physostigmine on impaired motor performance solely to sleep disruption.

We showed that the first sleep session after motor learning is crucial for memory consolidation, a finding in-line with other studies supporting the importance of first few hours of sleep post training (Huber R *et al*. 2006; Aeschbach D et al. 2008; Hanlon EC *et al*. 2009; Landsness EC et al. 2009; Landsness EC *et al*. 2011). We supported this conclusion by showing that physostigmine injections that increased ACh levels in the first few hours of sleep post training attenuated learning. We therefore conclude that reduced ACh levels during early sleep are crucial for motor memory consolidation. This finding is consistent with a previous human study related to the consolidation of declarative memory (Gais S and J Born 2004) although might not be directly comparable as the dosage used here was 10 times more. Thus, we propose that as for declarative memory, ACh levels might function as a switch, changing brain modes from the processes of memory encoding to the processes of memory consolidation (Hasselmo ME 1999; Hasselmo ME and J McGaughy 2004; Rasch BH *et al*. 2006).

We also asked whether further reducing ACh levels during early sleep would enhance or disrupt motor memory consolidation. Since, reducing ACh levels pharmacologically in wild type mice is a challenging manipulation, we used a chemogenetic approach utilizing CNO administered to transgenic mice to inactivate cholinergic neurons during early sleep (Anaclet C *et al*. 2015; Chen L et al. 2016; Kroeger D et al. 2017). Again, we did not measure ACh levels directly but inferred from behavior that levels were reduced. The rotarod speeds of mice on the Probe Day in the CNO group were significantly lower than those in the saline group, suggesting an effect of CNO injection. The seemingly negative effect of CNO might have adversely affected motor performance in comparison with Saline group (Gomez JL et al. 2017; Mahler SV and G Aston-Jones 2018; Manvich DF et al. 2018). There are mixed findings about whether selective ACh reduction affects motor performance (Conner JM et al. 2003; Gharbawie OA and IQ Whishaw 2003; Ramanathan D et al. 2009; Conner JM et al. 2010). Assuming that CNO had the intended action of reducing acetylcholine levels, we conclude that such acute manipulation adversely affects motor memory consolidation. These results support the idea that optimal acetylcholine levels in post-training sleep might be related to better memory consolidation. Increasing or decreasing these optimal ACh levels would disrupt motor memory consolidation.

Finally, we investigated whether it is muscarinic or nicotinic ACh receptors that play a crucial role in motor memory consolidation. Post-sleep motor performance was reduced by oxotremorine, a muscarinic acetylcholine receptor agonist, suggesting that activation of muscarinic receptors is undesirable for memory consolidation during early sleep. We did not perform EEG recordings after oxotremorine injection but assume that it would have reduced NREM sleep while increasing REM sleep as reported in previous studies with other muscarinic ACh receptor agonists e.g., pilocarpine and RS-86 (Lauriello J et al. 1993; Vanderwolf CH *et al*. 1993; Dringenberg HC and CH Vanderwolf 1996; Nissen C et al. 2006; Tejada S et al. 2007) or knockout mice with deleted genes for muscarinic receptors Chrm1 and Chrm3 (Niwa Y et al. 2018). Our results do not indicate the types of muscarinic receptors that might be involved in post-training memory consolidation but evidence obtained from Nissen et al (2006) showed that M1 receptor is not involved as its agonist (RS-86) altered sleep structure without altering declarative or procedural memory consolidation. There was no significant difference between oxotremorine and nicotine groups, suggesting that nicotinic receptors might also play some role in memory consolidation. Our results therefore support the idea that in natural early sleep, reduction in ACh on acetylcholine receptors is related to motor memory consolidation.

All drugs used in this study were injected systemically and would have acted on the peripheral as well as the central nervous system. Combinations of drugs could have been used to avoid peripheral effects of drugs e.g. physostigmine combined with methylatropine (Witkin JM 1989) to isolate central effects by reversing peripheral ones. The action of drugs on the peripheral nervous system therefore might also have contributed to changes in motor performance on the probe day. Nevertheless, since learning and memory formation for motor tasks primarily involves the central nervous systems, we argue that increase in ACh in the central nervous system prevented motor memory consolidation in experiments with physostigmine.

In our rotarod experiments, since the number of males and females was unequal in different groups, we were unable to determine whether there were Sex related effects on Probe Day performance or any interactions of Sex with Treatment or Training factors. Nevertheless, across all of our groups on the Training Day there was no effect of Sex on rotarod performance. Furthermore, we did not find any sex related effects on reach task performance when comparing Saline and Physostigmine groups. We therefore speculate that the role of cholinergic tone during early sleep in motor memory consolidation is similar in males and females. Although future experiments with Sex as a covariate could be undertaken, yet there have been no reports of sex-related performance differences on the rotarod or skilled reach tasks.

The findings of the present study are relevant to contemporary theories of how sleep contributes to motor memory consolidation. Our findings are consistent with the synaptic homeostatic theory of sleep that sleep facilitates memory consolidation by downscaling synaptic strengths to bring the synaptic load impacting each neuron to baseline levels, thus increasing signal to noise ratio of recently activated synapses (Tononi G and C Cirelli 2003). The homeostatic theory also suggests that low levels of ACh present during NREM sleep might mediate the synaptic downscaling process (Tononi G and C Cirelli 2006). Consistent with the homeostatic theory, when we increased ACh levels during early sleep, consolidation was impaired. Our findings are not inconsistent with the report that the cyclic succession of NREM and REM sleep is important for memory consolidation (Giuditta A *et al*. 1995; Ambrosini MV and A Giuditta 2001; Rasch and J Born 2013; Gais S and M Schonauer 2017). Both points of view suggest that NREM sleep causes selective weakening of non-adaptive memories (noise) and strengthening of adaptive (useful for survival) memories, whereas REM sleep is involved in the integration of adaptive memories into existing knowledge. In our study, physostigmine likely altered sleep structure disrupting the cyclic succession of NREM and REM sleep and thus causing impaired consolidation in this way. Alternatively, instead of disrupting the cyclic succession, physostigmine might have shifted NREM sleep to lighter phases, thus disrupting the strengthening of adaptive memories. The present results do not confirm a dual process theory that advocates that NREM sleep supports the consolidation of episodic memories while REM sleep facilitates the consolidation of motor memory (Stickgold R et al. 2000; Louie K and MA Wilson 2001; Mednick S et al. 2003; Peigneux P et al. 2003; Gais S and J Born 2004; Mascetti L et al. 2013). With respect to NREM sleep, our findings show that low ACh levels facilitate motor memory consolidation. Our findings are consistent with numerous other studies (see above) that do not support this theory. Perhaps the REM sleep dependent consolidation of motor memories is dependent on the type of task, but that assessment is beyond the scope of the studies described here (Chambers AM 2017). In parallel to the synaptic homeostasis hypothesis, our findings also agree with the active system consolidation hypothesis for hippocampus-dependent memories (HPM); in which low cholinergic tone during NREM sleep allows redistribution of HPM through reactivation while high cholinergic tone during REM sleep allows synaptic consolidation in neocortex (Diekelmann S and J Born 2010). Perhaps a similar two-stage model for motor memory involving cerebellum and motor cortex might allow consolidation of motor memories in the cortex (Krakauer JW and R Shadmehr 2006) and disruption of cholinergic tone during NREM sleep might impair the consolidation process as suggested by our results.

In conclusion, our results suggest that downregulated ACh levels during early NREM sleep are associated with motor memory consolidation. We used pharmacological manipulations for altering ACh levels, but future studies might use more selective methods. For example, by activating or silencing cholinergic neurons optogenetically with a feedback system to detect sleep states, one could clamp Ach levels in the desired brain region and study their role in memory consolidation using behavioral measures, electrophysiology, or optical imaging (Mohajerani MH et al. 2013). Furthermore, measurements to observe synaptic and biochemical changes during sleep would allow the study of structural correlates of memory consolidation.

## ACKNOWLEDGEMENTS

This study was supported by research fund provided by the Canadian Institutes of Health Research (CIHR) Grant# 390930, Natural Sciences and Engineering Research Council of Canada (NSERC) Discovery Grant #40352, Alberta Innovates (CAIP Chair) Grant #43568, Alberta Alzheimer Research Program Grant # PAZ15010 and PAZ17010, and Alzheimer Society of Canada Grant# 43674 to MHM; and NSERC Discovery Grant RGPIN-2017-03857 and NSF Grant 1631465 to BLM. We thank you Dr. Gerlinde Metz for providing us with the apparatus for performing the rotarod task. We thank Behroo Mirza Agha for her help with reach task training and data analyses. We thank Jessica Kuntz for her help with reach task data analyses. We thank Di Shao and the University of Lethbridge animal care staff for animal husbandry. Finally, we thank Drs. Masami Tatsuno and Bryan Kolb for useful discussions regarding the preparation of this manuscript.

## AUTHOR CONTRIBUTIONS

Q, SI, IQW, BLM and MHM designed the study. Q and SI performed all rotarod and reach task experiments. Q, SI, and IQW analyzed behavioral data. SI and SS collected and analyzed home cage video recording data. For electrophysiology experiments, MN performed animal surgeries for implanting electrodes, SI, Q, and MN collected data, and MN and SI analyzed data. MHM participated in all data analyses. SI, Q, IQW, and MHM wrote the manuscript, which all authors commented on and edited. MHM supervised the study.

## REFERENCES

Aeschbach D, Cutler AJ, Ronda JM. 2008. A role for non-rapid-eye-movement sleep homeostasis in perceptual learning. The Journal of neuroscience : the official journal of the Society for Neuroscience. 28:2766–2772.

Alger SE, Chambers AM, Cunningham T, Payne JD. 2015. The role of sleep in human declarative memory consolidation. Current topics in behavioral neurosciences. 25:269–306.

Ambrosini MV, Giuditta A. 2001. Learning and sleep: the sequential hypothesis. Sleep medicine reviews. 5:477–490.

Anaclet C, Pedersen NP, Ferrari LL, Venner A, Bass CE, Arrigoni E, Fuller PM. 2015. Basal forebrain control of wakefulness and cortical rhythms. Nature communications. 6:8744.

Aquilonius SM, Hartvig P. 1986. Clinical pharmacokinetics of cholinesterase inhibitors. Clinical pharmacokinetics. 11:236–249.

Boyce R, Glasgow SD, Williams S, Adamantidis A. 2016. Causal evidence for the role of REM sleep theta rhythm in contextual memory consolidation. Science. 352:812–816.

Brown DA. 2010. Muscarinic acetylcholine receptors (mAChRs) in the nervous system: some functions and mechanisms. Journal of molecular neuroscience : MN. 41:340–346.

Buhry L, Azizi AH, Cheng S. 2011. Reactivation, replay, and preplay: how it might all fit together. Neural plasticity. 2011:203462.

Buitrago MM, Ringer T, Schulz JB, Dichgans J, Luft AR. 2004. Characterization of motor skill and instrumental learning time scales in a skilled reaching task in rat. Behavioural brain research. 155:249–256.

Chambers AM. 2017. The role of sleep in cognitive processing: focusing on memory consolidation. Wiley interdisciplinary reviews Cognitive science. 8:1–14.

Chambers AM. 2017. The role of sleep in cognitive processing: focusing on memory consolidation. Wiley interdisciplinary reviews Cognitive science. 8.

Chen L, Yin D, Wang TX, Guo W, Dong H, Xu Q, Luo YJ, Cherasse Y, Lazarus M, Qiu ZL, Lu J, Qu WM, Huang ZL. 2016. Basal Forebrain Cholinergic Neurons Primarily Contribute to Inhibition of Electroencephalogram Delta Activity, Rather Than Inducing Behavioral Wakefulness in Mice. Neuropsychopharmacology. 41:2133–2146.

Clopath C. 2012. Synaptic consolidation: an approach to long-term learning. In. Cogn Neurodyn p 251–257.

Conner JM, Culberson A, Packowski C, Chiba AA, Tuszynski MH. 2003. Lesions of the Basal forebrain cholinergic system impair task acquisition and abolish cortical plasticity associated with motor skill learning. Neuron. 38:819–829.

Conner JM, Kulczycki M, Tuszynski MH. 2010. Unique contributions of distinct cholinergic projections to motor cortical plasticity and learning. Cerebral cortex (New York, NY : 1991). 20:2739–2748.

Dhingra D, Soni K. 2018. Behavioral and biochemical evidences for nootropic activity of boldine in young and aged mice. Biomedicine & pharmacotherapy = Biomedecine & pharmacotherapie. 97:895–904.

Diekelmann S, Born J. 2010. The memory function of sleep. Nat Rev Neurosci. 11:114–126.

Dringenberg HC, Vanderwolf CH. 1996. 5-Hydroxytryptamine (5-HT) agonists: effects on neocortical slow wave activity after combined muscarinic and serotonergic blockade. Brain research. 728:181–187.

Dringenberg HC, Vanderwolf CH. 1997. Neocortical activation: modulation by multiple pathways acting on central cholinergic and serotonergic systems. Experimental brain research. 116:160–174.

Dudai Y. 2004. The neurobiology of consolidations, or, how stable is the engram? Annual review of psychology. 55:51–86.

Dudai Y, Karni A, Born J. 2015. The Consolidation and Transformation of Memory. Neuron. 88:20–32.

Ebbinghaus H. 1885-Republished Translation 1964. Memory: A contribution to experimental psychology. Oxford, England: Dover.

Euston DR, Tatsuno M, McNaughton BL. 2007. Fast-forward playback of recent memory sequences in prefrontal cortex during sleep. Science. 318:1147–1150.

Frankland PW, Bontempi B. 2005. The organization of recent and remote memories. Nat Rev Neurosci. 6:119–130.

Gais S, Born J. 2004. Low acetylcholine during slow-wave sleep is critical for declarative memory consolidation. Proceedings of the National Academy of Sciences of the United States of America. 101:2140–2144.

Gais S, Plihal W, Wagner U, Born J. 2000. Early sleep triggers memory for early visual discrimination skills. Nature neuroscience. 3:1335–1339.

Gais S, Schonauer M. 2017. Untangling a Cholinergic Pathway from Wakefulness to Memory. Neuron. 94:696–698.

Genzel L, Kroes MC, Dresler M, Battaglia FP. 2014. Light sleep versus slow wave sleep in memory consolidation: a question of global versus local processes? Trends in neurosciences. 37:10–19.

Gharbawie OA, Whishaw IQ. 2003. Cholinergic and serotonergic neocortical projection lesions given singly or in combination cause only mild impairments on tests of skilled movement in rats: evaluation of a model of dementia. Brain research. 970:97–109.

Giuditta A, Ambrosini MV, Montagnese P, Mandile P, Cotugno M, Grassi Zucconi G, Vescia S. 1995. The sequential hypothesis of the function of sleep. Behavioural brain research. 69:157–166.

Gomez JL, Bonaventura J, Lesniak W, Mathews WB, Sysa-Shah P, Rodriguez LA, Ellis RJ, Richie CT, Harvey BK, Dannals RF, Pomper MG, Bonci A, Michaelides M. 2017. Chemogenetics revealed: DREADD occupancy and activation via converted clozapine. Science. 357:503–507.

Hanlon EC, Faraguna U, Vyazovskiy VV, Tononi G, Cirelli C. 2009. Effects of skilled training on sleep slow wave activity and cortical gene expression in the rat. Sleep. 32:719–729.

Hartvig P, Wiklund L, Lindstrom B. 1986. Pharmacokinetics of physostigmine after intravenous, intramuscular and subcutaneous administration in surgical patients. Acta anaesthesiologica Scandinavica. 30:177–182.

Hasselmo ME. 1999. Neuromodulation: acetylcholine and memory consolidation. Trends in cognitive sciences. 3:351–359.

Hasselmo ME, McGaughy J. 2004. High acetylcholine levels set circuit dynamics for attention and encoding and low acetylcholine levels set dynamics for consolidation. Progress in brain research. 145:207–231.

Huber R, Ghilardi MF, Massimini M, Ferrarelli F, Riedner BA, Peterson MJ, Tononi G. 2006. Arm immobilization causes cortical plastic changes and locally decreases sleep slow wave activity. Nature neuroscience. 9:1169–1176.

Huck SW, McLean RA. 1975. Using a repeated measures ANOVA to analyze the data from a pretest-posttest design: A potentially confusing task. Psychological Bulletin. 82:511–518.

Hut RA, Van der Zee EA. 2011. The cholinergic system, circadian rhythmicity, and time memory. Behavioural brain research. 221:466–480.

Jacobson TK, Howe MD, Schmidt B, Hinman JR, Escabi MA, Markus EJ. 2013. Hippocampal theta, gamma, and theta-gamma coupling: effects of aging, environmental change, and cholinergic activation. Journal of neurophysiology. 109:1852–1865.

Jafari-Sabet M, Jafari-Sabet AR, Dizaji-Ghadim A. 2016. Tramadol state-dependent memory: involvement of dorsal hippocampal muscarinic acetylcholine receptors. Behavioural pharmacology. 27:470–478.

Jendryka M, Palchaudhuri M, Ursu D, van der Veen B, Liss B, Katzel D, Nissen W, Pekcec A. 2019. Pharmacokinetic and pharmacodynamic actions of clozapine-N-oxide, clozapine, and compound 21 in DREADD-based chemogenetics in mice. Scientific reports. 9:4522.

Jenkins JG, Dallenbach KM. 1924. Obliviscence during Sleep and Waking. The American Journal of Psychology. 35:605–612.

Kali S, Dayan P. 2004. Off-line replay maintains declarative memories in a model of hippocampal-neocortical interactions. Nature neuroscience. 7:286–294.

Karni A, Meyer G, Rey-Hipolito C, Jezzard P, Adams MM, Turner R, Ungerleider LG. 1998. The acquisition of skilled motor performance: fast and slow experience-driven changes in primary motor cortex. Proceedings of the National Academy of Sciences of the United States of America. 95:861–868.

Klinzing JG, Kugler S, Soekadar SR, Rasch B, Born J, Diekelmann S. 2018. Odor cueing during slow-wave sleep benefits memory independently of low cholinergic tone. Psychopharmacology (Berl). 235:291–299.

Krakauer JW, Shadmehr R. 2006. Consolidation of motor memory. Trends in neurosciences. 29:58–64.

Kroeger D, Ferrari LL, Petit G, Mahoney CE, Fuller PM, Arrigoni E, Scammell TE. 2017. Cholinergic, Glutamatergic, and GABAergic Neurons of the Pedunculopontine Tegmental Nucleus Have Distinct Effects on Sleep/Wake Behavior in Mice. The Journal of neuroscience : the official journal of the Society for Neuroscience. 37:1352–1366.

Kudrimoti HS, Barnes CA, McNaughton BL. 1999. Reactivation of hippocampal cell assemblies: effects of behavioral state, experience, and EEG dynamics. The Journal of neuroscience : the official journal of the Society for Neuroscience. 19:4090–4101.

Landsness EC, Crupi D, Hulse BK, Peterson MJ, Huber R, Ansari H, Coen M, Cirelli C, Benca RM, Ghilardi MF, Tononi G. 2009. Sleep-dependent improvement in visuomotor learning: a causal role for slow waves. Sleep. 32:1273–1284.

Landsness EC, Ferrarelli F, Sarasso S, Goldstein MR, Riedner BA, Cirelli C, Perfetti B, Moisello C, Ghilardi MF, Tononi G. 2011. Electrophysiological traces of visuomotor learning and their renormalization after sleep. Clinical neurophysiology : official journal of the International Federation of Clinical Neurophysiology. 122:2418–2425.

Lauriello J, Kenny WM, Sutton L, Golshan S, Ruiz C, Kelsoe J, Rapaport M, Gillin JC. 1993. The cholinergic REM sleep induction test with pilocarpine in mildly depressed patients and normal controls. Biol Psychiatry. 33:33–39.

Li W, Ma L, Yang G, Gan WB. 2017. REM sleep selectively prunes and maintains new synapses in development and learning. Nature neuroscience. 20:427–437.

Louie K, Wilson MA. 2001. Temporally structured replay of awake hippocampal ensemble activity during rapid eye movement sleep. Neuron. 29:145–156.

MacLaren DA, Browne RW, Shaw JK, Krishnan Radhakrishnan S, Khare P, Espana RA, Clark SD. 2016. Clozapine N-Oxide Administration Produces Behavioral Effects in Long-Evans Rats: Implications for Designing DREADD Experiments. eNeuro. 3.

Mahler SV, Aston-Jones G. 2018. CNO Evil? Considerations for the Use of DREADDs in Behavioral Neuroscience. Neuropsychopharmacology. 43:934.

Manvich DF, Webster KA, Foster SL, Farrell MS, Ritchie JC, Porter JH, Weinshenker D. 2018. The DREADD agonist clozapine N-oxide (CNO) is reverse-metabolized to clozapine and produces clozapine-like interoceptive stimulus effects in rats and mice. Scientific reports. 8:3840.

Marchand M, Brossard P, Merdjan H, Lama N, Weitkunat R, Ludicke F. 2017. Nicotine Population Pharmacokinetics in Healthy Adult Smokers: A Retrospective Analysis. European journal of drug metabolism and pharmacokinetics. 42:943–954.

Mascetti L, Muto V, Matarazzo L, Foret A, Ziegler E, Albouy G, Sterpenich V, Schmidt C, Degueldre C, Leclercq Y, Phillips C, Luxen A, Vandewalle G, Vogels R, Maquet P, Balteau E. 2013. The impact of visual perceptual learning on sleep and local slow-wave initiation. The Journal of neuroscience : the official journal of the Society for Neuroscience. 33:3323–3331.

McCarthy EA, Maqsudlu A, Bass M, Georghiou S, Cherry JA, Baum MJ. 2017. DREADD-induced silencing of the medial amygdala reduces the preference for male pheromones and the expression of lordosis in estrous female mice. The European journal of neuroscience. 46:2035–2046.

McGaugh JL. 2000. Memory--a century of consolidation. Science. 287:248–251.

Mednick S, Nakayama K, Stickgold R. 2003. Sleep-dependent learning: a nap is as good as a night. Nature neuroscience. 6:697–698.

Miyamoto D, Hirai D, Murayama M. 2017. The Roles of Cortical Slow Waves in Synaptic Plasticity and Memory Consolidation. Front Neural Circuits. 11:92.

Mohajerani MH, Chan AW, Mohsenvand M, LeDue J, Liu R, McVea DA, Boyd JD, Wang YT, Reimers M, Murphy TH. 2013. Spontaneous cortical activity alternates between motifs defined by regional axonal projections. Nature neuroscience. 16:1426–1435.

Moyer TP, Charlson JR, Enger RJ, Dale LC, Ebbert JO, Schroeder DR, Hurt RD. 2002. Simultaneous analysis of nicotine, nicotine metabolites, and tobacco alkaloids in serum or urine by tandem mass spectrometry, with clinically relevant metabolic profiles. Clinical chemistry. 48:1460–1471.

Nagai H, de Vivo L, Bellesi M, Ghilardi MF, Tononi G, Cirelli C. 2017. Sleep Consolidates Motor Learning of Complex Movement Sequences in Mice. Sleep. 40:1–15.

Nissen C, Power AE, Nofzinger EA, Feige B, Voderholzer U, Kloepfer C, Waldheim B, Radosa MP, Berger M, Riemann D. 2006. M1 muscarinic acetylcholine receptor agonism alters sleep without affecting memory consolidation. Journal of cognitive neuroscience. 18:1799–1807.

Niwa Y, Kanda GN, Yamada RG, Shi S, Sunagawa GA, Ukai-Tadenuma M, Fujishima H, Matsumoto N, Masumoto KH, Nagano M, Kasukawa T, Galloway J, Perrin D, Shigeyoshi Y, Ukai H, Kiyonari H, Sumiyama K, Ueda HR. 2018. Muscarinic Acetylcholine Receptors Chrm1 and Chrm3 Are Essential for REM Sleep. Cell Rep. 24:2231–2247.e2237.

Olcese U, Bos JJ, Vinck M, Pennartz CMA. 2018. Functional determinants of enhanced and depressed interareal information flow in nonrapid eye movement sleep between neuronal ensembles in rat cortex and hippocampus. Sleep. 41:1–18.

Peigneux P, Laureys S, Fuchs S, Destrebecqz A, Collette F, Delbeuck X, Phillips C, Aerts J, Del Fiore G, Degueldre C, Luxen A, Cleeremans A, Maquet P. 2003. Learned material content and acquisition level modulate cerebral reactivation during posttraining rapid-eye-movements sleep. NeuroImage. 20:125–134.

Peters AJ, Chen SX, Komiyama T. 2014. Emergence of reproducible spatiotemporal activity during motor learning. Nature. 510:263–267.

Phillips K, Bartsch U, McCarthy A, Edgar D, Tricklebank M, Wafford K, Jones M. 2012. Decoupling of Sleep-Dependent Cortical and Hippocampal Interactions in a Neurodevelopmental Model of Schizophrenia. In. Neuron p 526–533.

Plihal W, Born J. 1997. Effects of early and late nocturnal sleep on declarative and procedural memory. Journal of cognitive neuroscience. 9:534–547.

Power AE, Vazdarjanova A, McGaugh JL. 2003. Muscarinic cholinergic influences in memory consolidation. Neurobiology of learning and memory. 80:178–193.

Pubchem. 2018. Physostigmine. In. Putcha L, Cintron NM, Tsui J, Vanderploeg JM, Kramer WG. 1989. Pharmacokinetics and oral bioavailability of scopolamine in normal subjects. Pharmaceutical research. 6:481–485.

Ramanathan D, Tuszynski MH, Conner JM. 2009. The basal forebrain cholinergic system is required specifically for behaviorally mediated cortical map plasticity. The Journal of neuroscience : the official journal of the Society for Neuroscience. 29:5992–6000.

Rasch, Born J. 2013. About sleep’s role in memory. Physiological reviews. 93:681–766.

Rasch BH, Born J, Gais S. 2006. Combined blockade of cholinergic receptors shifts the brain from stimulus encoding to memory consolidation. Journal of cognitive neuroscience. 18:793–802.

Saund J, Dautan D, Rostron C, Urcelay GP, Gerdjikov TV. 2017. Thalamic inputs to dorsomedial striatum are involved in inhibitory control: evidence from the five-choice serial reaction time task in rats. Psychopharmacology (Berl). 234:2399–2407.

Shiotsuki H, Yoshimi K, Shimo Y, Funayama M, Takamatsu Y, Ikeda K, Takahashi R, Kitazawa S, Hattori N. 2010. A rotarod test for evaluation of motor skill learning. Journal of neuroscience methods. 189:180–185.

Singh S, Bermudez-Contreras E, Nazari M, Sutherland RJ, Mohajerani M. 2018. Low-Cost Solution for Rodent Home-Cage Behaviour Monitoring. bioRxiv.

Sitaram N, Wyatt RJ, Dawson S, Gillin JC. 1976. REM sleep induction by physostigmine infusion during sleep. Science (New York, NY). 191:1281–1283.

Somani SM. 1989. Pharmacokinetics and pharmacodynamics of physostigmine in the rat after oral administration. Biopharmaceutics & drug disposition. 10:187–203.

Stachniak TJ, Ghosh A, Sternson SM. 2014. Chemogenetic synaptic silencing of neural circuits localizes a hypothalamus-->midbrain pathway for feeding behavior. Neuron. 82:797–808.

Stickgold R, Whidbee D, Schirmer B, Patel V, Hobson JA. 2000. Visual discrimination task improvement: A multi-step process occurring during sleep. Journal of cognitive neuroscience. 12:246–254.

Sutherland GR, McNaughton B. 2000. Memory trace reactivation in hippocampal and neocortical neuronal ensembles. Current opinion in neurobiology. 10:180–186.

Tejada S, Rial RV, Coenen AM, Gamundi A, Esteban S. 2007. Effects of pilocarpine on the cortical and hippocampal theta rhythm in different vigilance states in rats. The European journal of neuroscience. 26:199–206.

Tononi G, Cirelli C. 2003. Sleep and synaptic homeostasis: a hypothesis. Brain research bulletin. 62:143–150.

Tononi G, Cirelli C. 2006. Sleep function and synaptic homeostasis. Sleep medicine reviews. 10:49–62.

Tononi G, Cirelli C. 2014. Sleep and the price of plasticity: from synaptic and cellular homeostasis to memory consolidation and integration. Neuron. 81:12–34.

Vanderwolf CH, Raithby A, Snider M, Cristi C, Tanner C. 1993. Effects of some cholinergic agonists on neocortical slow wave activity in rats with basal forebrain lesions. Brain research bulletin. 31:515–521.

Vardy E, Robinson JE, Li C, Olsen RH, DiBerto JF, Giguere PM, Sassano FM, Huang XP, Zhu H, Urban DJ, White KL, Rittiner JE, Crowley NA, Pleil KE, Mazzone CM, Mosier PD, Song J, Kash TL, Malanga CJ, Krashes MJ, Roth BL. 2015. A New DREADD Facilitates the Multiplexed Chemogenetic Interrogation of Behavior. Neuron. 86:936–946.

Vyazovskiy VV, Olcese U, Lazimy YM, Faraguna U, Esser SK, Williams JC, Cirelli C, Tononi G. 2009. Cortical firing and sleep homeostasis. Neuron. 63:865–878.

Walker MP. 2009. The Role of Slow Wave Sleep in Memory Processing. J Clin Sleep Med. 5:S20–26.

Whishaw IQ. 2000. Loss of the innate cortical engram for action patterns used in skilled reaching and the development of behavioral compensation following motor cortex lesions in the rat. Neuropharmacology. 39:788–805.

Whishaw IQ, Faraji J, Kuntz J, Mirza Agha B, Patel M, Metz GAS, Mohajerani MH. 2017. Organization of the reach and grasp in head-fixed vs freely-moving mice provides support for multiple motor channel theory of neocortical organization. Experimental brain research. 235:1919–1932.

Whishaw IQ, Whishaw P, Gorny B. 2008. The structure of skilled forelimb reaching in the rat: a movement rating scale. Journal of visualized experiments : JoVE.

Wilson MA, McNaughton BL. 1994. Reactivation of hippocampal ensemble memories during sleep. Science. 265:676–679.

Witkin JM. 1989. Central and peripheral muscarinic actions of physostigmine and oxotremorine on avoidance responding of squirrel monkeys. Psychopharmacology (Berl). 97:376–382.

Yang G, Lai CS, Cichon J, Ma L, Li W, Gan WB. 2014. Sleep promotes branch-specific formation of dendritic spines after learning. Science. 344:1173–1178.

Yaroush R, Sullivan MJ, Ekstrand BR. 1971. Effect of sleep on memory. II. Differential effect of the first and second half of the night. Journal of experimental psychology. 88:361–366.

Young JM, Shytle RD, Sanberg PR, George TP. 2001. Mecamylamine: new therapeutic uses and toxicity/risk profile. Clinical therapeutics. 23:532–565.

Zhu H, Aryal DK, Olsen RH, Urban DJ, Swearingen A, Forbes S, Roth BL, Hochgeschwender U. 2016. Cre-dependent DREADD (Designer Receptors Exclusively Activated by Designer Drugs) mice. Genesis (New York, NY : 2000). 54:439–446.

