## Supplementary Figures for "Low acetylcholine during early sleep is important for motor memory consolidation"

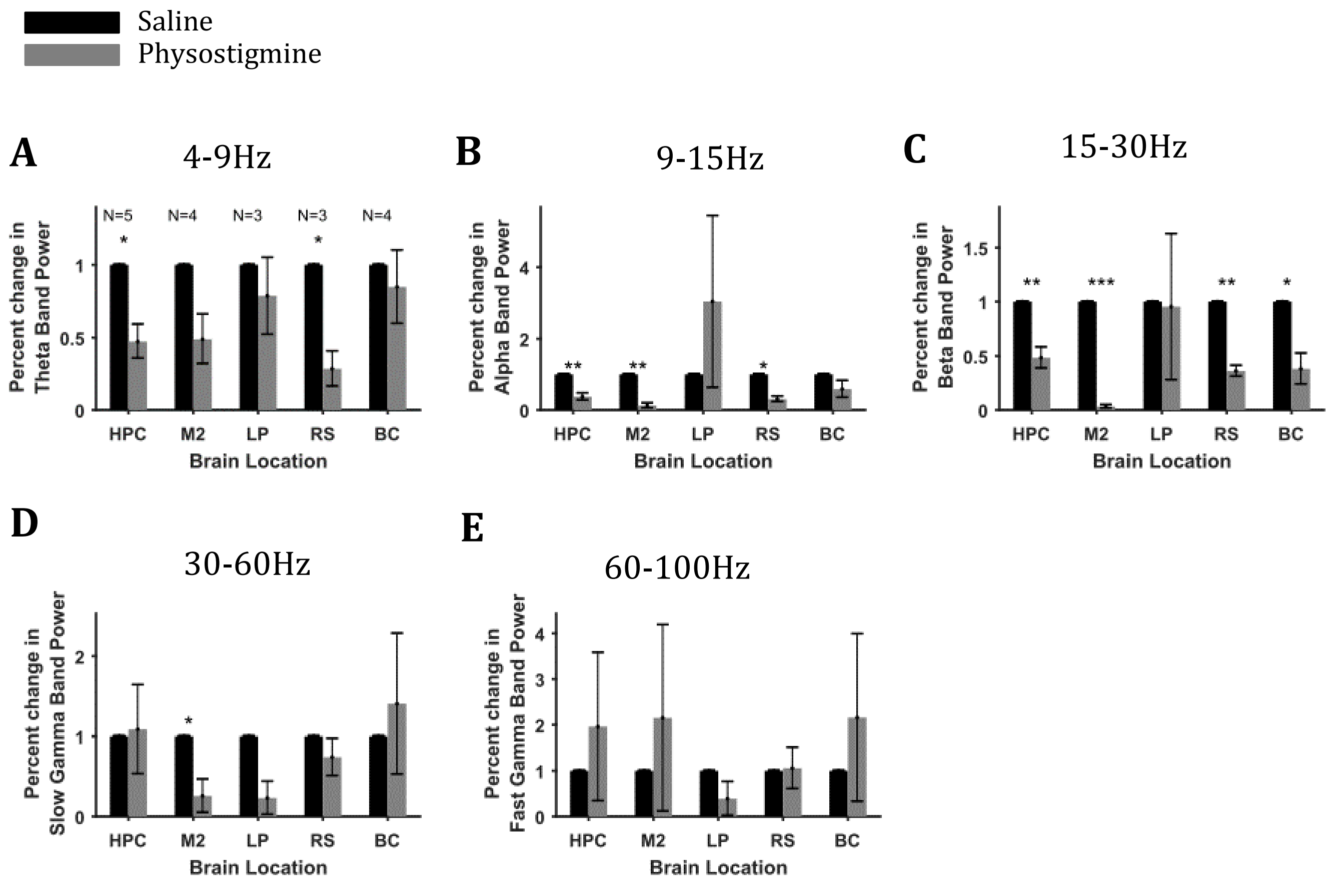


**Figure S1. Sleep characterization using electrophysiology in the first hour after the onset of NREM sleep. (A-E)** Percentage change in power after physostigmine injections in higher frequency bands within the first hour after the onset of NREM sleep. Error bars represent SEM. N represents number of animals in A and is same for B-E.

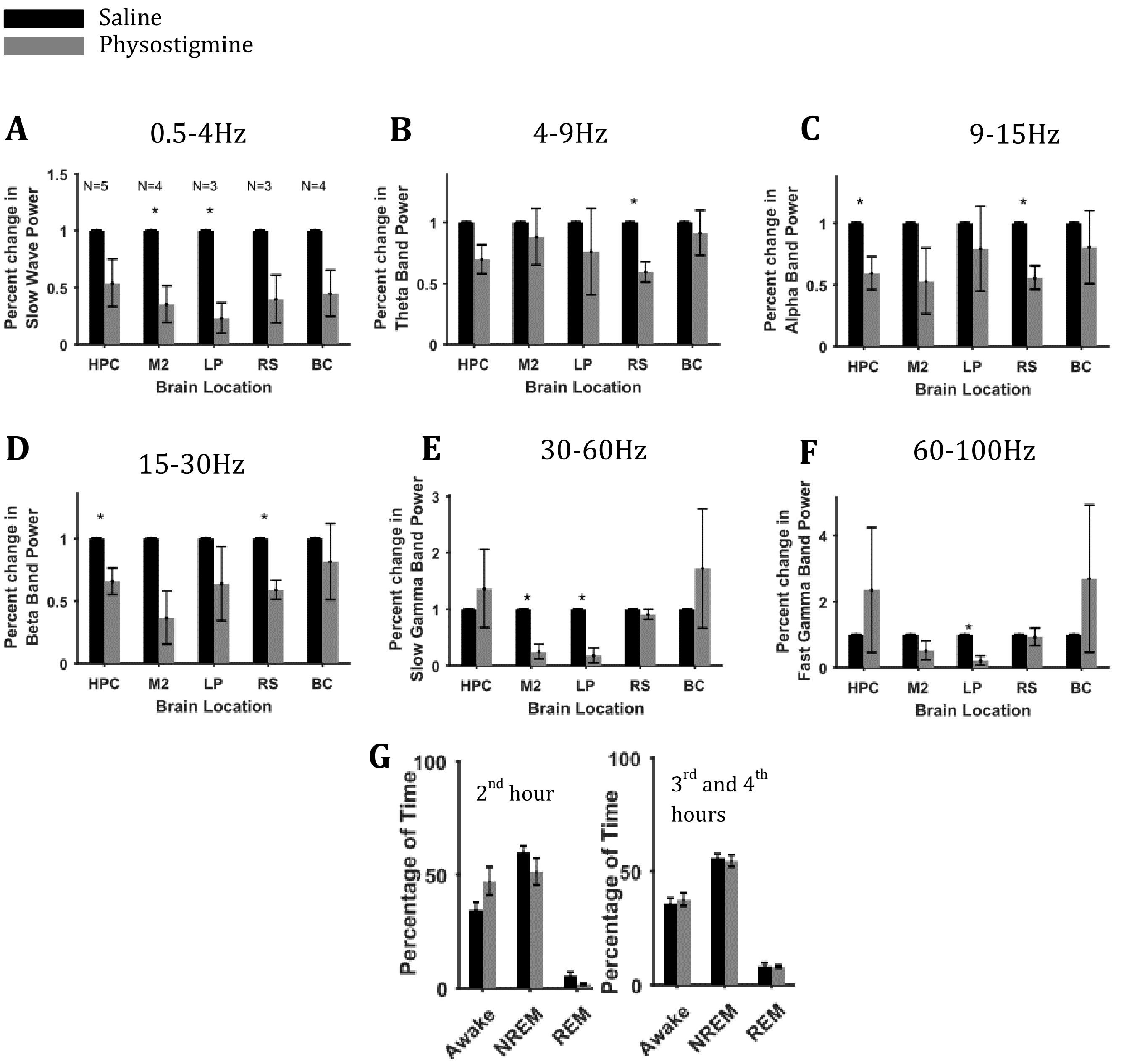


**Figure S2. Sleep characterization using electrophysiology of 2nd and 3rd hours after the onset of NREM sleep. (A-F)** Percentage change in power in different frequency bands after physostigmine injections, during two hours duration, one hour after the onset of NREM sleep i.e. in the 2nd and 3rd hours after the onset of NREM sleep. Error bars represent SEM. N represents number of animals shown in A and is same for C-F. **(G)** Percentage of time for different states in the 2nd and 3-4th hours (N=5).
